# Early blood-spinal cord barrier stabilization preserves peri-lesional tissue and improves recovery after SCI

**DOI:** 10.64898/2025.12.03.692136

**Authors:** Chen Chen, Yan Sun, Xiaolong Du, Seung-Young Lee, Xiaoting Wang, Qi Han, Wenhui Xiong, Xiangbing Wu, Yiping Zhang, Christopher B. Shields, Xiaoming Jin, Ji-Xin Cheng, Wei Wu

## Abstract

Vascular disruption is an early and critical event in spinal cord injury (SCI), but its contribution remains poorly understood. Using novel in vivo two-photon dual-dye imaging, we found simultaneous blood-spinal cord barrier (BSCB) leakage and venous dilation in both the injury epicenter and adjacent transitional segment after cervical SCI in rats. Notably, vascular permeability in the transitional zone preceded axonal and neuronal loss, revealing a therapeutic window. Systemic delivery of ferulic acid-glycol chitosan (FA-GC) nanoparticles, a membrane-sealant, rapidly stabilized the compromised vasculature, reduced neuronal loss in the transitional region, and improved forelimb muscle strength. These findings identified acute vascular leakage beyond the injury epicenter as a driver of secondary pathology and highlight early vascular stabilization as promising therapeutic strategy.

## Introduction

Spinal cord injury (SCI) triggers a cascade of pathological process that extend beyond the primary mechanical insult, ultimately resulting in irreversible neurological deficits. Among the earliest and most consequential of these secondary events is the disruption of blood-spinal cord barrier (BSCB), a specialized vascular interface that maintains homeostasis by tightly regulating molecular and cellular exchange between the circulatory system and spinal cord parenchyma (*1*). This barrier is composed of endothelia cells sealed by tight junctions, supported by pericytes, basement membrane, and astrocytic end-feet (*2*). Unlike the more restrictive blood-brain barrier (BBB), the BSCB exhibits inherently higher permeability and lower tight junction protein expression, rendering it particularly vulnerable to trauma-induced breakdown (*3, 4*).

Following SCI, rapid and sustained BSCB leakage permits the extravasation of serum proteins, immune cells, and inflammatory mediators into spinal cord tissue (*5, 6*). These changes exacerbate edema, oxidative stress, and cell death, accelerating lesion expansion and impeding repair (*7–9*). Despite recognition of BSCB dysfunction as hallmark of SCI, most studies have relied on static, postmortem assessment of vascular damage, limiting insight into its temporal dynamics and spatial heterogeneity (*7, 10, 11*). It therefor remains unclear whether vascular disruption merely a consequence of tissue injury or an active driver of secondary degeneration (*12–14*).

To address this fundamental question, we applied high-resolution *in vivo* two-photon imaging with dual-dye vascular labeling (*11*) to visualize the evolution of BSCB permeability after cervical contusion SCI in rats. This approach allowed us to quantify vascular leakage and venous dilation across anatomically distinct spinal segments – specifically the injury epicenter (C7), a peri-lesional “transitional” segment (C6), and remote uninjured region (C5). Surprisingly, we observed that BSCB breakdown in the transitional zone occurred concurrently with that at epicenter, despite the absence of overt mechanical damage. Vascular leakage in this segment preceded axonal degeneration, oligodendrocyte loss, and neuronal death by several hours to days, identifying it as an early and spatially distinct site of vulnerability.

Given the temporal window between BSCB leakage and subsequent cellular degeneration, we hypothesized that early restoration of vascular integrity could prevent or reduce secondary damage in the transitional zone. We therefore tested the efficacy of our self-synthesized ferulic acid-glycol chitosan (FA-GC) nanoparticles (*15*), a membrane-sealing nanotherapy that selectively target compromised vasculature and stabilizes disrupted endothelial membranes. FA-GC combines the anti-oxidative and anti-inflammatory properties of ferulic acid with the biocompatibility and self-assembly capabilities of glycol chitosan, enabling selective accumulation at sites of injury-induced membrane damage.

We demonstrate that systemic delivery of FA-GC shortly after SCI rapidly attenuated BSCB leakage in the transitional segment, preserved ventral horn neurons, and improved forelimb motor strength in the acute recovery phase. These effects occurred without direct targeting the epicenter, underscoring the functional relevance of early intervention in peri-lesional tissue.

Our findings establish that acute, region-specific vascular disruption beyond the lesion epicenter is not merely a bystander phenomenon but a therapeutic target. Furthermore, the FA-GC nanoparticles effect as a minimally invasive strategy for membrane repair and neuroprotection. By integrating dynamic vascular imaging with targeted nanotherapy, this study provides new mechanistic insight and therapeutic direction for early intervention after SCI.

## Results

### Acute vascular dilation and leakage occur at the injury epicenter and transitional regions following SCI

To investigate the spatiotemporal dynamics of acute spinal vascular changes following SCI, we labeled spinal blood vessels with sequentially fluorescent tracers and track them using *in vivo* two-photon imaging. We assessed vascular responses across three cervical segments – the injury epicenter (C7), transitional region(C6), and remote region (C5) – via an imaging window spanning C5-C7 after laminectomy (Fig. 1a). Two key parameters were measured: vascular permeability and vessel diameter, reflecting barrier breakdown and hemodynamic disruption. Vascular permeability was quantified by comparing the fluorescent intensity of the rhodamine dextran tracer (Rho, red, 70kDa) outside versus inside individual vessels at pre– and post-SCI time points respectively (Fig. 1b-e, teal and purple bars), with higher ratio indicating greater vascular permeability. In sham controls, the tracer remained confined within vessels, showing a dark background across all 3 segments (Fig. 1b, f-h, lilac lines). However, within 0.5 hour (h) post-injury, significant leakage emerged in both the epicenter and transitional segment, while the remote segment remained unaffected (Fig. 1c, Supplemental Fig. 1a, b). In leaking vessels, histological analysis confirmed structural disruption of the vascular wall (Supplementary c-d”). In sham cords, RECA-1+ endothelial cells formed smooth, continuous linings closesly apposed to laminin-labeled basement membranes, with evenly distributed nuclei and clear lumens, indicating intact barrier integrity and normal vessel tone. In contrast, after SCI, endothelial and basement membrane layers, and distorted or collapsed lumens, signifying endothelial detachment, basement membrane rupture, and loss of vascular tone.

**Figure 1.**
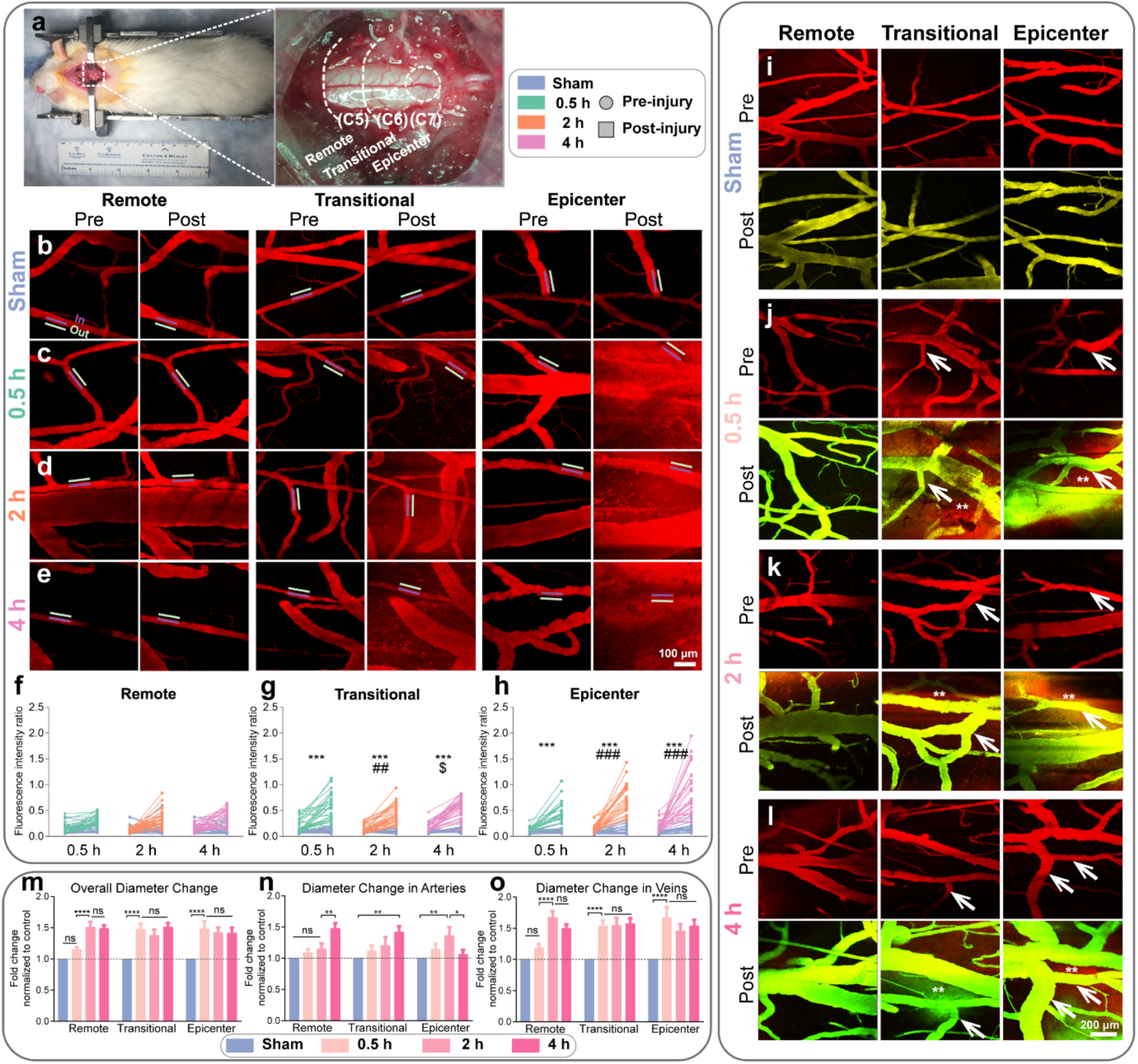
Acute vascular dilation and leakage occur simultaneously in both the epicenter and transitional region after a midline C7 contusive SCI. **(a)** Images of the observed cervical spinal cord segments during *in vivo* imaging under the surgical microscope: C5 (remote), C6 (transitional), and C7 (epicenter) after a C7 midline contusion. Vascular permeability of the same vessels pre– and post-SCI, measured via rhodamine-dextran(Rho, red, 70 kDa) fluorescence intensity (purple bar: inside vessel; white bar: outside vessel), is shown for (b) Sham, (c) 0.5 hour (h), (d) 2 h, and (e) 4 h post-injury. Higher outside/inside ratio indicates greater permeability. Quantification of the spatiotemporal changes of vessel permeability. Quantified spatiotemporal changes in permeability for **(f)** remote, **(g)** transitional, and **(h)** epicenter regions. n = 3-4 animals per group; for each region per animal, n = 8-10 blood vessel segments. Repeated measure two-way ANOVA followed by LSD post hoc test. *: versus sham control. #: versus 0.5 h post-injury. $: versus 2 h post-injury. * # $ P < 0.05; ** ## $$ P < 0.01; *** ### $$$ P < 0.001; **** #### $$$$ P < 0.0001. Each line represents pre– and post-injury changes in one blood vessel segment. The measurements of vessel diameter before and after SCI are also shown in **(i)** Sham, **(j)** 0.5 h, **(k)** 2 h, and **(l)** 4 h post-injury. Quantification of the spatiotemporal changes of diameters in **(m)** overall vessels, **(n)** arteries, and **(o)** veins in sham, 0.5 h, 2 h and 4 h post-injury. n = 8-10 animals per group; for each region per animal, n = 5-7 blood vessel segments. For each segment, one-way ANOVA followed by Tukey’s multiple comparison tests were performed. ns, not significant; * P < 0.05; ** P < 0.01; *** P < 0.001; **** P <0.0001 vs sham control. Data were presented as mean ± s.e.m.

To assess vessel changes, we administered a second tracer, Fluorescein isothiocyanate dextran (FITC, green, 70kDa), at 0.5 h, 2 h, and 4 h post-injury following an initial pre-injury Rho (*11*). Sham controls exhibited stable vessel diameters across all segments (Fig. 1i). In contrast, by 0.5 h post-SCI, marked vasodilation (white arrows) and blood content extravasation (**) occurred almost simultaneously in epicenter and transitional segments, with no significant change in the remote segment (Fig. 1j). Vessel diameter increased markedly by approximately a 1.5-fold in the epicenter and transitional segment (Transitional: 1.472±0.664, Epicenter: 1.481±0.609, p<0.0001, compared with sham; Fig.1m). Dilation in the remote segment emerged later, at 2 hours, and persisted across all segments through 4 hours (Fig. 1k, l). Further analysis classified vessels into arteries and veins based on established criteria, revealing that venous dilation was the earliest and prominent contributor to vessel diameter increases, with arterial responses emerging later and more modestly (Fig. 1n, o). Collectively, these findings demonstrate that acute vascular leakage and dilation occur simultaneously in the epicenter and adjacent transitional region, extending beyond the injury core following SCI. This hemodynamic dysfunction likely contributes to vascular congestion and accelerates barrier breakdown, identifying vascular instability as a initiator of secondary injury in peri-lesional tissue.

### Vascular Disruption Affects Both White and Gray Matter Tissue Sections

To explore whether the vascular changes observed by *in vivo* imaging occur in a deeper parenchyma and persist over time, we conducted immunofluorescence staining for immunoglobulin G (IgG) and laminin (Table. 1) to detect barrier disruption in the white and gray matter of the C5-7 segments up to 7 days (d) post-injury. IgG, which does not cross blood-brain barrier (BBB) and blood-spinal cord barrier (BSCB), appears as diffuse parenchymal staining when barrier integrity is compromised, allowing quantification of permeability changes. In the white matter, we focused on the dorsal column, which is directly impacted by the midline contusive SCI (Fig. 2a). Consistent with our *in vivo* imaging results, significant IgG-positive vessel disruption occurred simultaneously in the epicenter (78.23±14.54%, p<0.0001) and the transitional segment (51.76±16.06%, p=0.0003) as early as 0.5 h post-injury, while the remote segment showed no notable leakage (24.01±25.13%, p=0.4215; Fig. 2b-c). IgG extravasation intensified at 2 and 4 hours, and by 1 d post-injury, vessel leakage became significant in the remote segment (44.63±18.91%, p=0.0434). This disruption persisted across all three regions through 7 days (Remote: 52.2±7.279%, p=0.0015, Transitional: 52.86±7.608%, p=0.0005, Epicenter: 56.33±14.74%, p<0.0001; versus sham).

**Figure 2.**
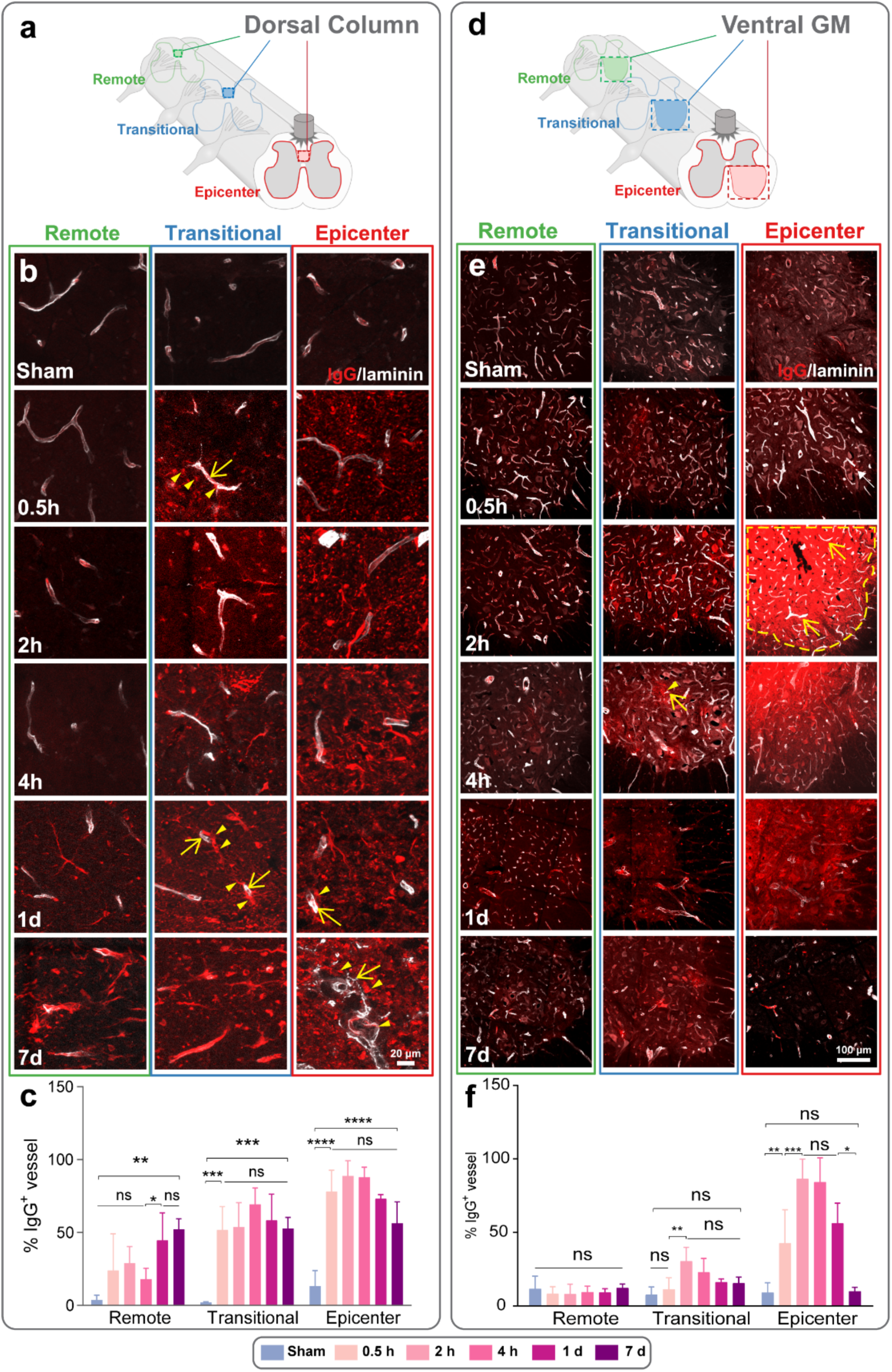
Histology shows acute vascular leakage in both the epicenter and transitional regions after a midline C7 contusive SCI. **(a)** A schematic diagram shows sampled regions in the dorsal column white matter. **(b)** In the dorsal column white matter, vessels become leaky, labeled by immunoglobulin G (IgG) and laminin, in both the epicenter and transitional region starting 0.5 hours (h) post-injury. The vessels were monitored at various time points up to 7 days (d) post-injury. Yellow arrows indicate the leaky vessels. Yellow arrowheads point at IgG leaking outside the vessels. **(c)** Quantification of the percentage of IgG^+^ vessels over total laminin^+^ vessels in the dorsal column. **(d)** A schematic diagram shows the observed regions in the ventral part of gray matter. **(e)** In the ventral gray matter, vessels also become leaky in the epicenter starting at 0.5 h post-injury and the transitional segment starting at 2 h post-injury. The vessels were monitored at different time points up to 7 d post-injury. Yellow arrows indicate the leaky vessels. Yellow dashed lines show the area IgG was leaked. **(f)** Quantification of the percentage of IgG+ vessels over total laminin^+^ vessels in the ventral gray matter. For **(c)** and **(f)**, n = 7-8 for sham, 0.5 h, 2 h, and 4 h post-injury. n = 6 for 1 d and 7 d post-injury. For each segment, one-way ANOVA followed by Tukey’s multiple comparison tests were performed. * P < 0.05; ** P < 0.01; *** P < 0.001; **** P<0.0001. All data were presented as mean ± s.d.

In the gray matter, we examined the ventral region, which contains motor neurons critical for motor outputs as well as diverse interneuron populations, and is known to be particularly vulnerable to degeneration post SCI (Fig. 2d). IgG leakage was evident in the epicenter as early as 0.5 hours, but not in the transitional or remote segments (Fig. 2e, f). The transitional segment exhibited delayed leakage starting at 2 hours, which progressed alongside the epicenter through 4 hours. By 1day, significant IgG leakage persisted only in the epicenter, and by 7 days, it was no longer detectable in either the epicenter or transitional segment. The remote segment showed no significant leakage throughout.

Together, These findings demonstrate two key points: (1) histological evidence validates the simultaneous leakage in the dorsal column of epicenter and transitional segment observed by *in vivo* imaging; and (2) acute vessel leakage occur in both the white and gray matter following acute SCI but follows distinct spatiotemporal dynamics depending on region and tissue type.

### Delayed Axonal and Myelin Loss in the Transitional Segment Dorsal Column

The dorsal column, comprising ascending and descending axonal tracts, received direct impact from the midline contusive injury. To evaluate the acute axonal response, we examined axons and myelination in the epicenter (C7), transitional (C6), and remote (C5) segments (Fig. 3a). In sham controls, in cross-sectional views showed consistent numbers of myelinated axons (green, SMI-31) across three segments, with myelin (red, MBP) tightly ensheathing axons to enhance action potential conduction (Fig. 3b). At 0.5 hours post-injury, axonal loss was detected in the epicenter, but absent in the transitional and remote segments (Fig. 3c). Myelin sheath loosened around the axons, significantly increasing total myelin area and myelin area per axon in both transitional and remote segments as early as 0.5 hours (Fig. 3d, Supplemental Fig. 2a), consistent with prior reports (*16, 17*), while in the Epicenter, significant myelin area increase was started at 2 hours post injury. At 0.5, 2, and 4 hours, axonal deterioration progressed only in the epicenter, with persistent myelin detachment reflected by elevated myelin area metrics. At 1 day post-injury, axonal loss extended to transitional and remote segments, accompanied by reduced myelin area, due to fragmentation of the dismantled myelin and rapid clearance of myelin debris by activated macrophages and microglia (*18–21*). By 7 days, significant axon and myelin loss persisted across all three segments (Fig. 3d). Notably, while myelin changes occurred immediately, axonal loss in the transitional segment was delayed relative to the earlier vascular disruption observed in this region.

**Figure 3.**
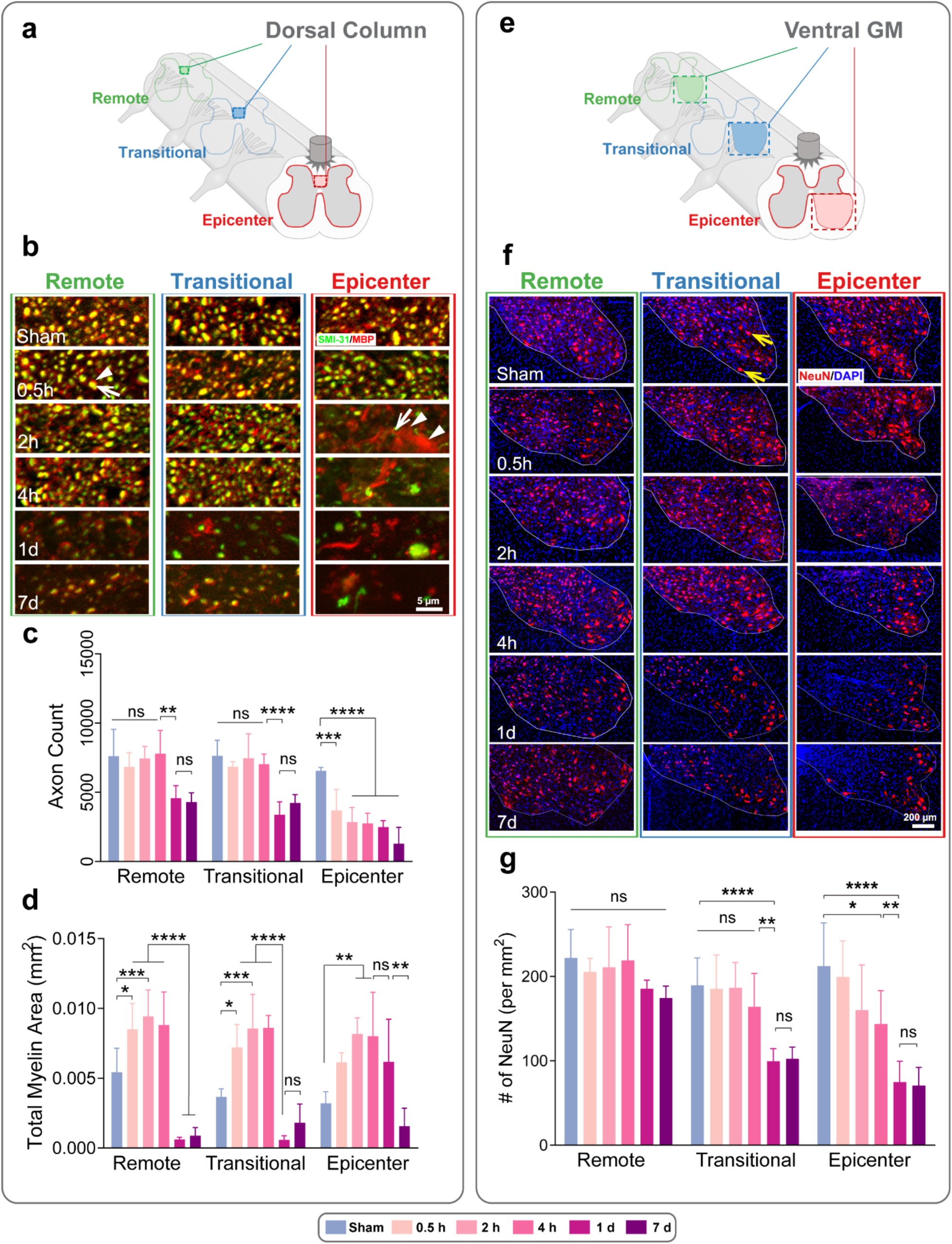
Spatiotemporal loss of axons, myelin, and neurons after a midline C7 contusive SCI. **(a)** A schematic diagram shows the sampled regions in the dorsal column white matter for axons and myelin. **(b)** Axon and myelin are double-labeled by SMI-31 (green) and MBP (red) and are examined in the remote, transition, and epicenter regions in Sham, 0.5 hours (h), 2 h, 4 h, 1 day (d), and 7 d post-injury. White arrows indicate axons and white arrowheads indicate myelin in a loose form. **(c)** Quantification of axon number in measured area of dorsal column. **(d)** Quantification of total myelin area in measured area of dorsal column. **(e)** A schematic diagram shows the sampled regions in the ventral gray matter. **(f)** Neurons are double labeled with NeuN and DAPI, and their densities are measured in 3 different regions at different time points up to 7 d post-injury. In the epicenter, the neuronal loss started at 4 h post-injury; in the transitional region, the neuronal loss started at 1 d post-injury which lags behind vessel leakages. Yellow arrows indicate neurons. **(g)** Quantification of neuronal density in the ventral gray matter. For **(c)**, **(d),** and **(g)**, n = 7-8 for sham, 0.5 h, 2 h, and 4 h post-injury. n = 6 for 1 d and 7 d post-injury. For each segment, one-way ANOVA followed by Tukey’s multiple comparison tests were performed. * P < 0.05; ** P < 0.01; *** P < 0.001; **** P <0.0001. All data were presented as mean ± s.d.

These findings demonstrate that axonal degeneration in the transitional zone is delayed relative to vascular disruption, with myelin changes preceding structural axonal loss.While the epicenter exhibited rapid axonal deterioration within hours of injury, the transitional and remote regions initially showed only myelin loosening, reflected by enlarged myelin area measurements. Significant axonal loss in these segments emerged only after one day, coindiding with myelin fragmentation and clearance by activated macrophages. Thus, vascular dysfunction and myelin instability represent early events, whereas axonal degeneration in the transitional region follows a more protracted cours, highlighting a temporal dissociation between vascular injury, myelin pathology, and axonal loss.

### Delayed Neuronal Loss in the Transitional Segment Ventral Horn

We next examined neuronal loss in the ventral gray matter (Fig. 3e). In sham controls, intact neurons, labeled by NeuN (red) and DAPI (blue), exhibited a consistent density of approximately 200 per mm^2^ across all 3 segments (Fig. 3f-g). After injury, neuronal density in the remote segment (C5) remained stable through 7 days. In the epicenter (C7), density dropped significantly to 143.50±39.61 per mm^2^ by 4 hours (p=0.0343, versus sham), and further to 70.58±21.55 per mm^2^ by 7 days (p<0.0001, versus sham). In the transitional segment (C6), neuronal density remained unchanged until 1day post-injury, when it decreased to 99.38±15.03 per mm^2^ (p=0.0001, versus sham), stabilizing at 102.30±13.91 per mm^2^ by 7 days after injury (p=0.0002, versus sham). This decline in the transitional segment occurred after the earlier vascular disruption observed in the same region. Quantitatively, total neurons counts mirrored these density trends (Supplemental Fig. 2b).

Analysis of neuronal density in the ventral horn revealed a segment-specific and delayed pattern of loss compared with dorsal column. Neurons in the epicenter were highly vulnerable, showing significant reduction as early as 4 hours, with progressive decline through 7 days. Bye contrast, the transitional segment maintained stable neuronal density until one day post-injury, at which point a sharp decline occurred that persisted through 7 days. The remote segment remained unaffected throughout. These findings highlight a one-day lag between vascular disruption and neuronal degeneration in the transitional gray matter, underscoring the role of early vascular instability as a precursor to later neuronal loss.

### Time– and Region-Specific Glial Responses to SCI

Microglia, critical for central nervous system (CNS) surveillance and secondary injury after SCI (*22–25*), were assessed for activation in the ventral gray matter using Iba1 staining (Fig. 4a). In sham controls, the resident microglia displayed a ramified morphology across all 3 segments (Fig. 4b, yellow arrow). Post-injury, microglia shift to a hypertrophied and round shape in response to trauma (Fig. 4b, yellow arrowhead) (*26, 27*). At 0.5 hours after injury, the epicenter showed fewer ramified microglia (p=0.0069) without an increase in hypertrophied type (p=0.9367, Fig. 4b-c). This decline intensified at 2 and 4 hours (p<0.0001, versus sham), with ramified microglia nearly absent by 4 hours, suggesting microglial loss (p<0.0001). In contrast, the transitional and remote segments maintained stable microglia counts through 4 hours. By 1 day, hypertrophied microglia increased in the epicenter and transitional segments (Fig. 4b, yellow arrowhead; Fig. 4c, p=0.0032), with ramified forms remaining low (p=0.0013). At 7 days, the total number of microglia was drastically increased across all three segments (p<0.0001, Fig. 4c).

**Figure 4.**
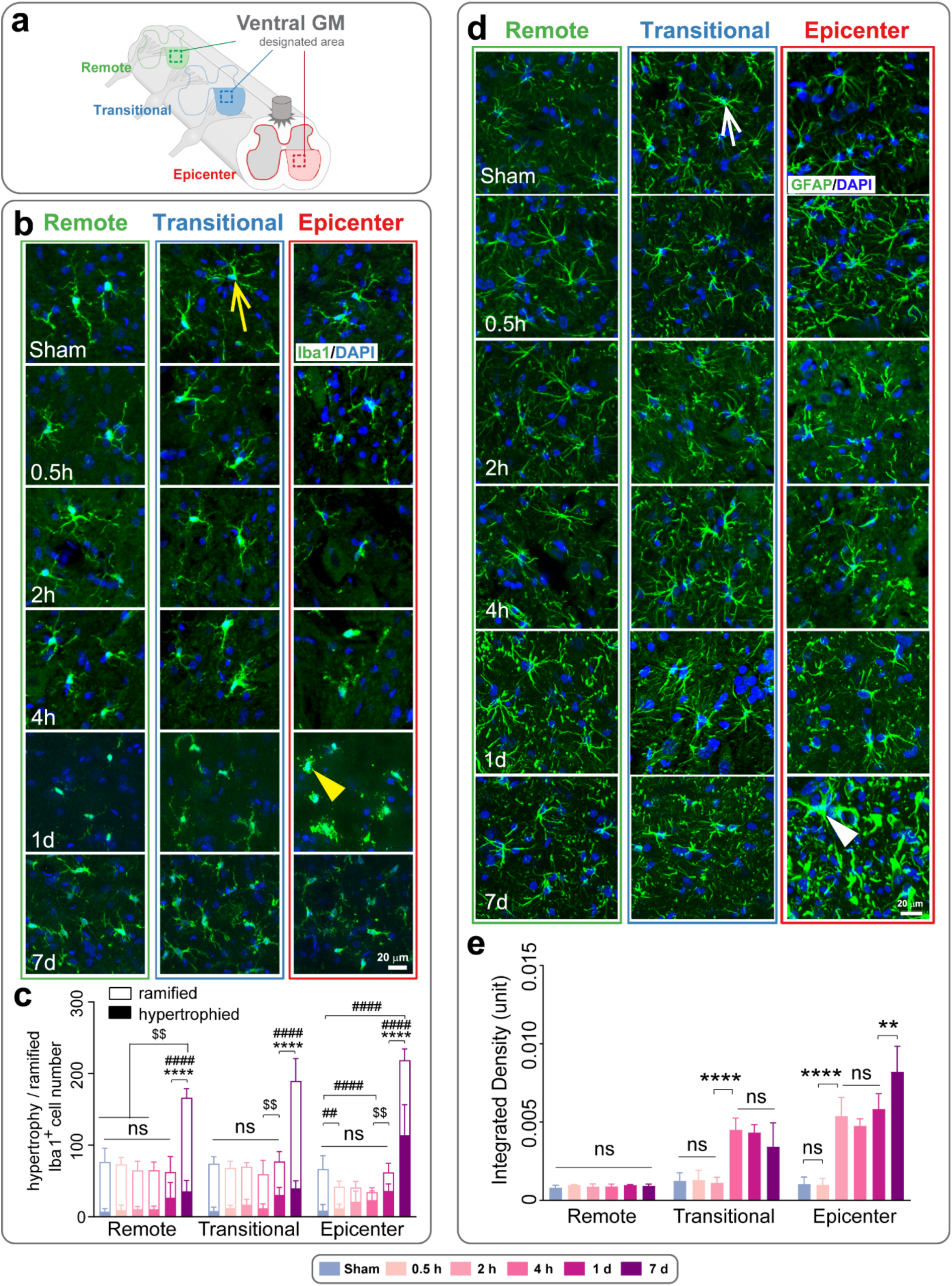
Time and region-specific activation of microglia and astrocytes after a C7 midline contusive SCI. **(a)** A schematic diagram shows the sampled regions in the ventral gray matter. **(b)** Microglial activation is examined using immunofluorescent double-labeling of Iba1 and DAPI in 3 different spinal regions at different time points up to 7 days. Yellow arrow indicates a ramified microglia and yellow arrowhead indicates a hypertrophied microglia. **(c)** The number of hypertrophied and ramified microglia was quantified and displayed in stacked histogram. n = 6 per group. For each segment, one-way ANOVA followed by Tukey’s multiple comparison tests were performed. *: Total cell number. $: Hypertrophied cell number. #: Ramified cell number. ns, not significant; * # $ P < 0.05; ** ## $$ P < 0.01; *** ### $$$ P < 0.001; **** #### $$$$ P < 0.0001. **(d)** Representative images of the ventral gray matter shows the activation of astrocytes, immunofluorescent double-labeled with GFAP and DAPI, at different regions and different time points. White arrow indicates a normal astrocyte and white arrowhead indicates a reactive astrocyte. **(e)** Quantification of the integrated density of GFAP expression in the designated area of ventral gray matter. n = 7-8 for sham, 0.5 hour (h), 2 h, and 4 h post-injury. n = 6 for 1 d and 7 d post-injury. For each segment, one-way ANOVA followed by Tukey’s multiple comparison tests were performed. ns, not significant; * P < 0.05; ** P < 0.01; *** P < 0.001; **** P <0.0001. All data were presented as mean ± s.d.

To evaluate the time– and region-specific astrocytic responses to SCI, the cell density and glial fibrillary acid protein (GFAP) expression levels were examined in the same region (Fig. 4a). Sham controls showed protoplasmic and stellate astrocytes (Fig. 4d, sham, white arrow). At 0.5 hours post-SCI, no changes in either GFAP expression or cell density was observed across all 3 segments (Fig. 4d-e, Supplemental Fig. 3b). By 2 hours, astrocytes retracted their processes, and meanwhile GFAP expression was elevated in the epicenter (Fig. 4d-e). Similar elevated GFAP expressions were observed in the transitional segment, while the remote segment remained constant as sham controls. At the time point, changes in cell density were not detectable in all 3 segments (Supplemental Fig. 3b). At 4 h after injury, fewer astrocytes were observed and GFAP remained at a higher level in the epicenter (Fig. 4e). The transitional segment also showed an elevated level of GFAP expression yet no changes in cell density. The remote segment was not affected in both cell density and GFAP expression. At 1 day, epicenter astrocyte density recovered (Supplemental Fig. 3b), with GFAP remaining elevated in both epicenter and transitional segments, while the remote segment remained at the same level as sham controls. No changes in cell density were detected in the transitional nor remote segment. At 7 days, the epicenter showed a marked increase of cell density and GFAP expression (Fig. 4d, white arrowhead), while the transitional segment sustained high GFAP expression without density changes (Supplemental Fig. 3b), lagging the vascular disruption demonstrated earlier (Fig. 4e).

Oligodendrocytes response were quantified using CC1 staining in both the dorsal column and ventral gray matter across 3 segments (Fig. 5a, d). In the dorsal column, the epicenter quickly lost oligodendrocyte density at 0.5 hours (p<0.0001), while transitional and remote segments remained stable until 1 day post-injury (p=0.0003 & p<0.0001 respectively; Fig. 5b, c). A slight recovery was observed in the epicenter by day 7. In the ventral gray matter, all segments maintained the cell density until 1 day, when loss emerged (p<0.0001), persisting through 7 days (Fig. 5e, f). In both white and gray matter, oligodendrocyte loss in the transitional segment followed vascular leakage.

**Figure 5.**
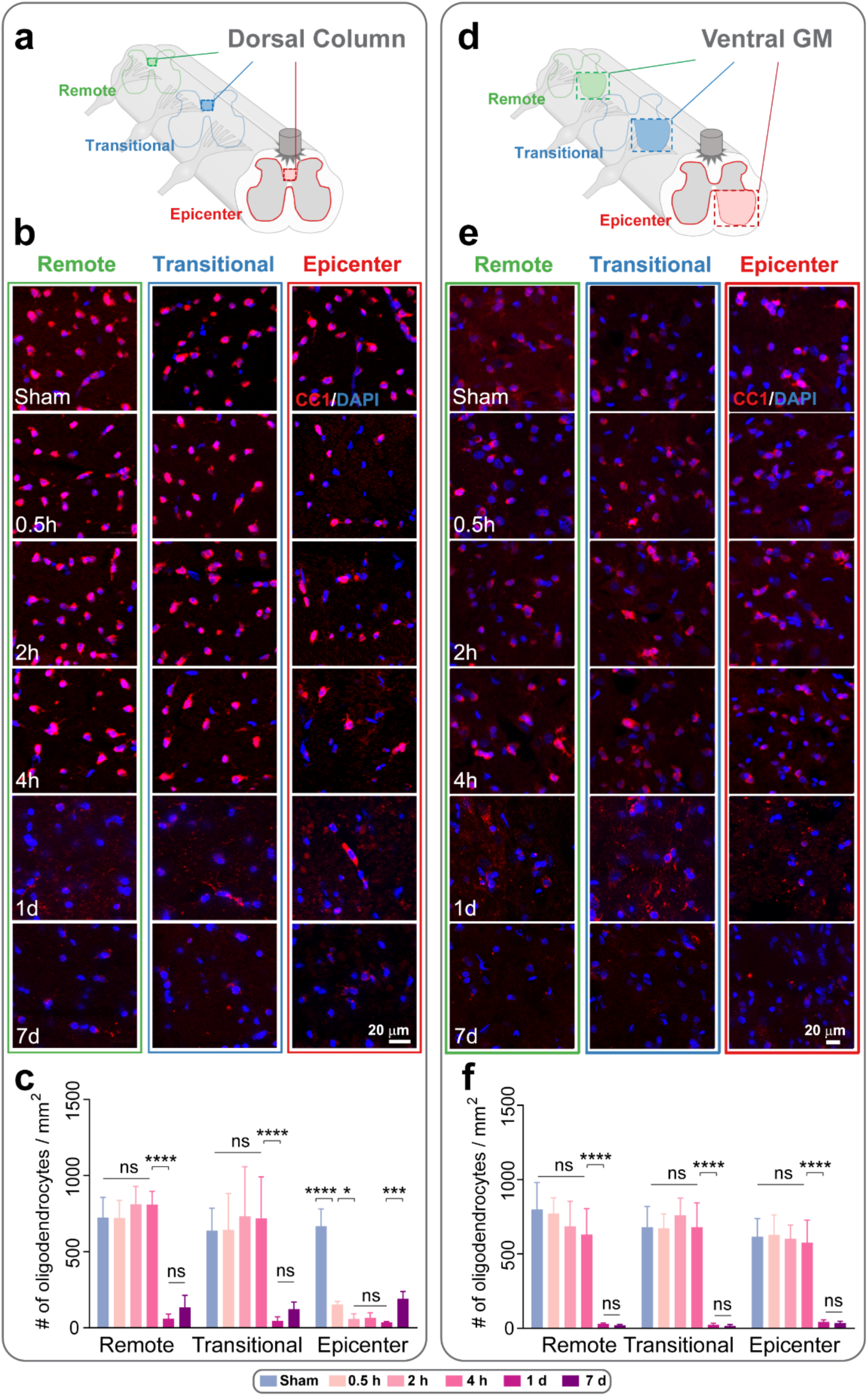
Loss of oligodendrocytes in both the dorsal column and ventral gray matter after a C7 midline contusive SCI. **(a)** A schematic diagram shows the sampled regions in the dorsal column white matter. **(b)** Oligodendrocytes are immunofluorescent double-labeled with CC1 and DAPI and are visualized in 3 different spinal regions at different time points. **(c)** The density of oligodendrocytes is calculated in the defined region. **(d)** A schematic diagram shows the sampled regions in the ventral gray matter. **(e)** Representative images show CC1-positive oligodendrocytes in the ventral gray matter at different regions and at different time points. **(f)** Quantification of oligodendrocyte density in the ventral gray matter. For **(c)** and **(f)**, n = 7-8 for sham, 0.5 hour (h), 2 h, and 4 h post-injury. n = 6 for 1 day (d) and 7 d post-injury. For each segment, one-way ANOVA followed by Tukey’s multiple comparison tests were performed. ns, not significant; * P < 0.05; ** P < 0.01; *** P < 0.001; **** P <0.0001. All data were presented as mean ± s.d.

Together, these results reveal a sequential glial response following SCI. Microglia rapidly transformed from ramified to hypertrophied forms, showing early loss in the epicenter followed by proliferation and widespread acrtivation by 7 days. Astrocytes exhibited delyed but sustained activation, with elevated GFAP expression beginging at 2 hours and progressing to increased density and glial scar formation in the epicenter and transitional regions. Ologodendrocyte loss occurred earliest in the epicenter and leter in the adjacent segments, closely raralleling the pattern of vascular disruption and contributing to to secondary demyelination.

### FA-GC Nanoparticles Seal Disrupted BSCB and Protect Transitional Segment Ventral Horn Neurons

Given the time window between early vascular dysfunction and delayed neuronal damage, the increased BSCB permeability in the transitional segment may be reversible, offering a target for neuroprotection. We tested self-synthesized ferulic acid-glycol chitosan (FA-GC) nanoparticles, known for their membrane-sealing properties (*15*), to address this. *in vivo* two-photon imaging at 2 hours post-sci showed that intravenous FA-GC significantly reduced rhodamine dextran (Rho) tracer extravasation into the parenchyma across all three segments (Fig. 6a-b, d). However, FA-GC did not affect the SCI-induced vascular dilation (Fig. 6c).

**Figure 6.**
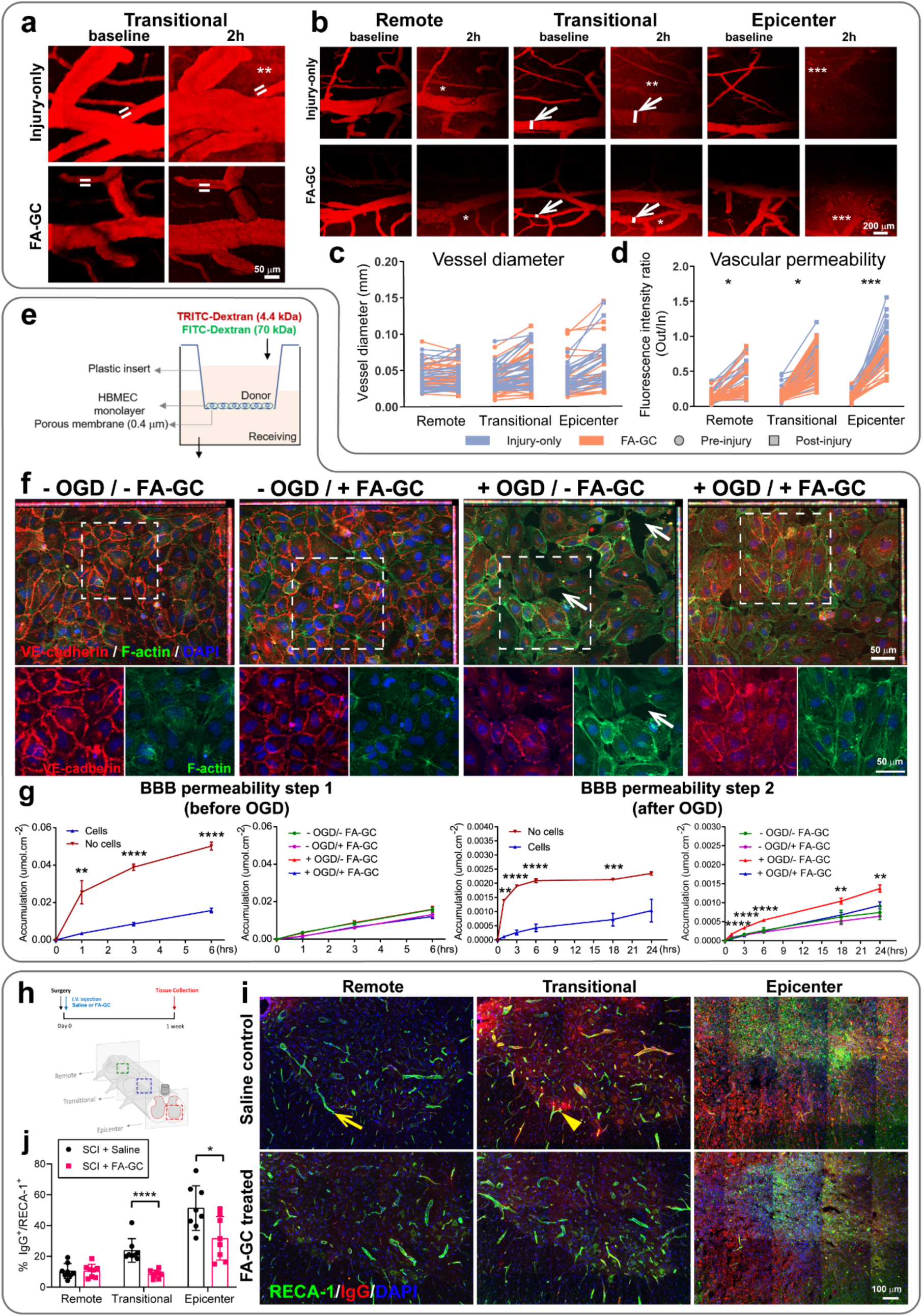
FA-GC nanoparticles effectively seal the vascular membrane and reduce vascular permeability. **(a)** FA-GC treatment reduced the vascular permeability at 2 hours (h) post-injury comparing to injury-only group. The white bars indicate the location where measures the vascular permeability, and ** indicates the leaking area. **(b)** Intravital two-photon images show changes in vessel diameter and vascular permeability after FA-GC treatment comparing to injury-only controls. White arrows indicate the measured blood vessels, and white bars indicate the diameters of the blood vessels. * indicates the leaking area, and the severity of leakage is shown by increased number of symbols. **(c)** The changes in vessel diameter, impacted by both the injury and FA-GC treatment, were shown by measuring the same blood vessels at pre– and post-injury timepoints across three regions (epicenter, transitional, and remote). **(d)** The changes in vascular permeability were calculated by measuring the ratio of fluorescence intensity of rhodamine-dextran (70kDa) outside versus inside. For **(c)** and **(d)**, n = 5 for injury-only, n = 9 for FA-GC; for each region per animal, n = 6-8 blood vessel segments. Repeated measure two-way ANOVA followed by LSD post hoc test. * P < 0.05; ** P < 0.01; *** P < 0.001; **** P < 0.0001. Each line represents pre– and post-injury changes in one blood vessel segment. **(e)** Schematic drawing of the *in vitro* BBB model. Full details of this experiment are provided in the methods section and supplementary figures. **(f)** Under the conditions of control or 2 hours (h) of oxygen glucose deprivation (OGD), human brain microvascular endothelial cells (HBMECs) were immediately treated with FA-GC or medium control and stained at 24 h after OGD for F-actin^+^ stress fibres (green) and the adhesion junctional protein VE-cadherin (red), and counterstained with DAPI for nuclear labeling. OGD-induced stress fibre formation was significantly attenuated by FA-GC treatment. White dashed rectangular frames indicate the location of the enlarged areas listed below.White arrows indicate the enlarged gap between endothelial cells. **(g)** The diffusion coefficient of the two different-size fluorescent tracers was quantified by accumulation of the tracers in the receiving chamber. The TRITC-dextran tracer (4.4 kDa) was measured at 0-6 h after control non-OGD conditions to confirm the integrity of HBMEC monolayer. The FITC-dextran tracer (70 kDa) was quantified at 0-24 h after OGD or control non-OGD conditions, with FA-GC treatment or medium control. Data represent four independent experiments. Repeated measure two-way ANOVA followed by LSD post hoc test. * P < 0.05; ** P < 0.01; *** P < 0.001; **** P <0.0001 vs control or – OGD/-FA-GC. Data were presented as mean ± s.e.m. **(h)** Illustration of the timeline and observed locations of FA-GC *in vivo* experiments. **(i)** Representative images of the ventral gray matter cross sections with FA-GC treatment or saline control after SCI were stained with endothelial cell marker RECA-1 (green) and IgG (red), and counterstained with DAPI for nuclear labeling. Yellow arrow indicates a normal blood vessel with no IgG leakage, and yellow arrowhead points at a IgG-leaked blood vessel. **(j)** The percentage of IgG^+^/RECA-1^+^ blood vessel segments over all RECA-1^+^ blood vessels was quantified at 3 different spinal segments with FA-GC treatment or saline control. N = 8 per group. For each segment, comparison of means between two groups was accomplished by the Student’s t-test (two-tailed). * P < 0.05; ** P < 0.01; *** P < 0.001; **** P <0.0001. Data were presented as mean ± s.d.

To confirm FA-GC’s sealing effect on CNS barriers, we implemented an *in vitro* BBB model with human brain microvascular endothelial cells (HBMECs; Fig. 6e, Supplemental Fig. 4a, b), subjected to 2 hours of oxygen glucose-deprivation (OGD). Previous studies indicated that OGD induced EC actin cytoskeletal rearrangement by promoting the stress fiber formation of F-actin from G-actin (*28*). Consistent with the previous findings, robust stress fiber formation was visualized by immunostaining of F-actin 24 hours after OGD (Fig. 6f, – OGD/-FA-GC, + OGD/-FA-GC). Meanwhile, wide gaps were formed between ECs under the OGD condition (white arrows; Supplemental Fig. 4c). FA-GC treatment attenuated stress fiber formation, reducing F-actin staining (Fig. 6f). Permeability was assessed in two steps: first, a 4.4kDa TRITC-dextran tracer confirmed the HBMEC monolayer integrity up to 6 h (Supplemental Fig. 4a, b). Four groups of cells grew under the same condition and showed the same level of integrity pre-OGD(Fig. 6g, step 1); second, a 70kDa FITC-dextran tracer revealed OGD-induced hyperpermeability from 1 to 24 hours (Fig. 6g, step 2). BBB hyperpermeability emerged at 1 h post-OGD with increased diffusion of 70kDa tracer across the EC barrier and were persistent up to 24 h after OGD. FA-GC effectively reduced the accumulation of tracers at 1-24 h after OGD (Fig. 6g, BBB permeability step 2). At 1 week after SCI, we also assessed the level of IgG leakage after FA-GC application by quantifying the percentage of IgG^+^/RECA-1^+^ vessel segments in the ventral gray matter (Fig. 6h). FA-GC suppressed IgG leakage in both epicenter and transitional segment (Fig. 6i, j). These data collectively support that FA-GC could seal the disrupted BBB and reduce extravasation of blood contents.

To evaluate neuroprotection of FA-GC, we quantified ventral horn neurons at 1 week nanoparticles following SCI (Fig. 7a). Since the cell type, size, and density varies in the gray matter depending on different laminae, counting the total neuronal number may mask the regional differences. Based on the anatomical structure of cervical spinal cord, the gray matter was divided into 3 regions: dorsal horn (DH), intermediate zone (IZ), and ventral horn (VH) (Fig. 7b) (*29*). No significant difference was detected with FA-GC treatment in all 3 segments compared with saline controls (Fig. 7c). In the ventral horn, however, FA-GC treatment saved more neurons in the transitional segment compared with saline controls (Fig. 7d; p=0.0116). These findings demonstrate that FA-GC seals disrupted BSCB and mitigates neuronal loss in the transitional segment.

**Figure 7.**
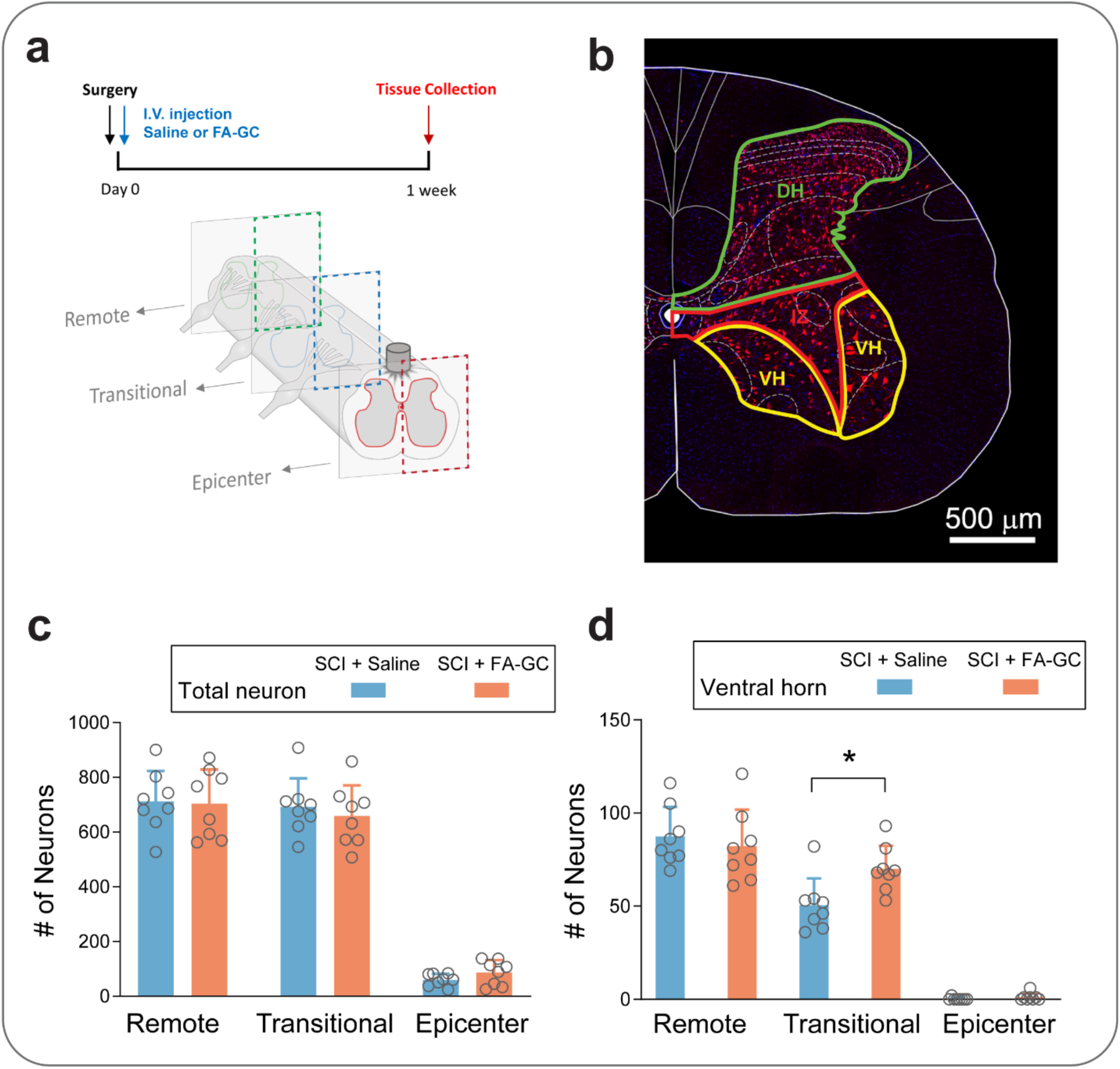
FA-GC nanoparticles rescue ventral horn neurons in the transitional region after a midline C7 contusive SCI. (**a**) Illustration of timelines and sampling of tissue after the C7 contusive spinal cord injury (SCI) and FA-GC or saline treatment. **(b)** Representative image of NeuN^+^/DAPI^+^ neurons in the locations where were quantified. DH, dorsal horn. IZ, intermediate zone. VH, ventral horn. **(c)** The number of neurons in half of the gray matter was quantified across three regions with FA-GC or saline administration. **(d)** The number of neurons in the ventral horn was quantified across 3 spinal segments with FA-GC or saline administration. For **(c)** and **(d)**, n = 8 per group. For each segment, comparison of means between two groups was accomplished by the Student’s t-test (two-tailed). * P < 0.05. Data were presented as mean ± s.d.

### FA-GC Nanoparticles Enhance Forelimb Muscle Strength

To evaluate the functional impact of the midline C7 contusive SCI, we assessed motor functions for 6 weeks (Fig. 8a). Grip strength were selected to measure forelimb muscular strength after FA-GC treatment (Fig. 8a-b). Tests were conducted every other day alternately in the first week, then weekly from weeks 2 to 6 post-SCI. The injury group exhibited persistent forelimb strength deficits in grip strength from 2 to 42 days post-SCI compared to sham controls, despite no body weight differences (Fig. 8d). FA-GC treatment significantly improved forelimb strength from 14 to 6 weeks compared to saline controls (Fig. 8c). This functional recovery aligns with the preservation of ventral horn neurons by FA-GC nanoparticles, suggesting a link between neuronal rescue and motor recovery.

**Figure 8.**
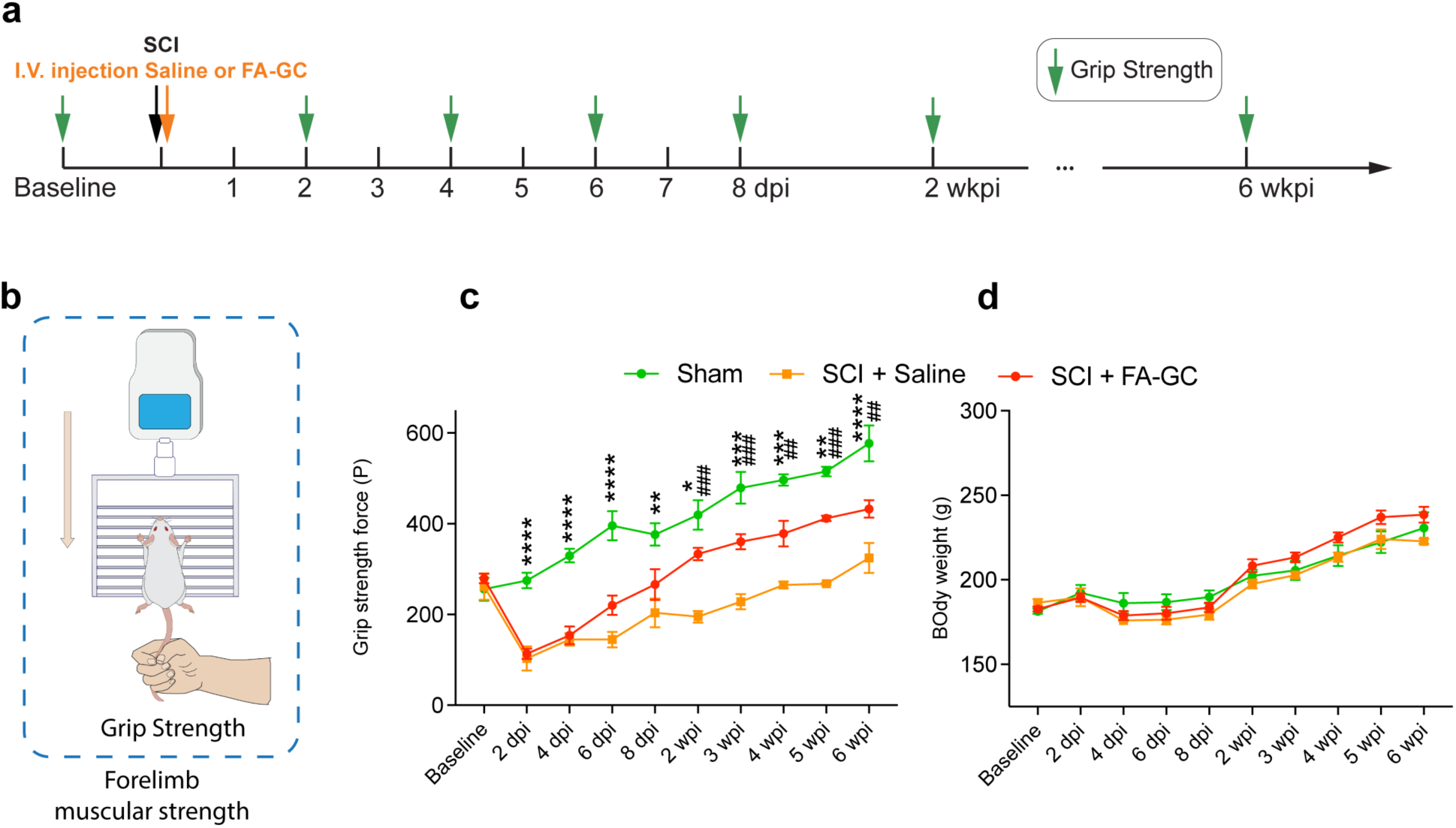
FA-GC nanoparticles improve forelimb muscle strength after a midline C7 contusive SCI. (**a**) The timeline of grip strength test on Sham, SCI rats after FA-GC treatment or saline control was evaluated before and at 0-6 wk after SCI. **(b)** Illustration of grip strength test. **(c)** The grip strength force in FA-GC treated groups was significantly strengthened as compared to the saline group starting at 14 d after injury. **(d)** The changes in body weight were no different among Sham, FA-GC treated and saline-treated groups during the same time. n = 4 for Sham, n = 3 for SCI + Saline, and n = 8 for SCI + FA-GC. At each time point, comparison of means between two groups was accomplished by the Student’s t-test (two-tailed).*: Sham vs FA-GC; #: FA-GC VS Saline. #* P < 0.05; ##** P < 0.01; ###*** P < 0.001; ####**** P <0.0001. Data were presented as mean ± s.d.

## Discussion

The vascular interface between blood and neural tissue is essential for spinal cord homeostasis and is rapidly compromised following traumatic injury (*30*). Here, we reveal that disruption of the BSCB occurs not only at the injury epicenter but also in a previously underappreciated adjacent region, defined here as the transitional segment. Using *in vivo* dual-dye two-photon imaging, we demonstrate that vascular leakage and venous dilation emerge in this segment within hours of injury, preceding axonal and neuronal loss (Supplemental Fig. 5). These findings indicate that early microvascular dysfunction is not restricted to the lesion core and may initiate secondary injury in neighboring tissue.

Previous studies have documented BSCB breakdown in postmortem tissue after SCI, but lacked the spatiotemporal resolution to determine when and where permeability changes first arise (*7, 9, 31*). By directly visualizing the BSCB leakage across three spinal levels, our results identify a spatially defined and temporally dynamic injury gradient. Importantly, the transitional segment exhibits delayed cellular degeneration relative to the epicenter, yet shows early vascular leakage, suggesting a therapeutic window in which barrier stabilization could prevent irreversible damage (*32, 33*). Although the epicenter displays more severe IgG leakage, particularly in the ventral horn (Fig. 2), multiple factors, including direct mechanical insult, intense inflammation, and oxidative stress, contribute to its complex pathology, making it difficult to isolate the role of vascular permeability alone(*9, 34*). In contrast, the transitional zone presents a relatively simpler pathophysiological context where early vascular disruption precedes overt neuronal loss. This spatial separation between vessel damage and neuronal degeneration offers a unique opportunity to test the hypothesis that microvascular leakage initiates secondary injury. Additionally, damage within the epicenter is often irreversible by the time of intervention, whereas protecting the transitional segment may help preserve viable tissue and improve functional outcomes (*35*). Thus, the transitional zone provides both a mechanistic and therapeutic rationale for targeted vascular stabilization strategies.

To test this concept, we administered FA-GC nanoparticles. In prior work, FA-GC improved locomotor recovery and reduced tissue damage after SCI(*15, 36*). In the present work, we reveal a mechanistic basis for this protection: FA-GC acts primarily on the transitional segment, a region previously overlooked but now identified as a site of early vascular vulnerability. FA-GC reduced BSCB leakage, preserved ventral horn neurons, and improved motor function. These findings emphasize the importance of both timing and spatial precision in vascular repair strategies and establish FA-GC not only as a general neuroprotectant but also as a targeted intervention capable of intercepting secondary damage driven by acute vascular instability.

The composition of the leaked content from compromised vasculature is not benign. For example, fibrinogen – a major blood protein that enters the CNS after BSCB disruption – has been shown to exacerbate injury by activating TGF-β signaling, promoting scar formation, and enhancing neuroinflammation (*3, 14, 37*). In addition to fibrinogen, other blood-derived factors such as thrombin, complement proteins, albumin, and hemoglobin breakdown products can activate microglia and astrocytes, induce oxidative stress, and potentiate excitotoxicity, thereby amplifying secondary degeneration of neurons and oligodendrocytes (*3, 38–43*). Thus, once the BSCB is breached, the spinal parenchyma is exposed to a sustained “toxic spill” of plasma proteins and inflammatory mediators that not only mark the injury but also actively propagate it. Preventing or rapidly limiting the entry of these components is therefore expected to reduce glial activation, attenuate scar formation, and preserve myelin and neuronal integrity. Our findings suggest that vascular leakage is not merely a biomarker of injury but also a driver of pathophysiological cascades, further justifying interventions aimed at early vascular sealing.

In summary, this study identifies a spatially distinct and therapeutically accessible region of acute vascular dysfunction following SCI. Our results demonstrate that early systemic administration of FA-GC nanoparticles selectively stabilizes the BSCB, reduces neuronal loss, and improves motor function. These findings establish the transitional segment as a critical therapeutic target and introduce vascular sealing as a viable early intervention strategy for spinal cord injury.

## Materials and Methods

### Animals

A total of 128 female Sprague-Dawley rats (6 weeks old, 180-220 gram) were purchased from Envigo RMS, Inc. (Indianapolis, IN). They were housed 2 per cage with an air filter in an environmentally controlled facility with a 12-hour/12-hour light/dark cycle. Food and water were fully accessible to the animals throughout the entire study. All procedures were approved by the Institutional Animal Care and Use Committee (IACUC) of Indiana University School of Medicine and Institutional Biosafety Committee (IBC) and strictly followed the National Institutes of Health (NIH) Guide on humane care and the use of laboratory animals.

In the first study, a total of 76 rats were used and randomly assigned into 2 groups of experiments: *in vivo* spinal vascular imaging (*n* = 32), immunofluorescence labeling (*n* = 44). In the second study, a total of 52 rats were used and randomly assigned into 3 groups of experiments: *in vivo* spinal vascular imaging (*n* = 18), immunofluorescence labeling (*n* = 16), and forelimb functional assessments (*n* = 18).

### Spinal cord injury

Animals were assigned for two independent studies. In preparation for surgery, the rats were injected with ketamine (87.7mg/kg) and xylazine (12.3mg/kg) mixture intraperitoneally to induce deep anesthesia. In the first study, a total of 32 rats in the *in vivo* spinal vascular imaging experiment received a laminectomy from the 5^th^ to 7^th^ cervical vertebrae; another 44 rats in the histological experiment received a laminectomy at the 7^th^ cervical vertebrae. In the second study, a total of 18 rats in the in vivo spinal vascular imaging experiment received a laminectomy from the 5^th^ to 7^th^ cervical vertebrae; both 16 rats in the histological experiment and 18 rats in the behavioral experiment received a laminectomy at the 7^th^ cervical vertebrae. For all SCI performed in this study, the animals received a mild-to-moderate contusive injury at the midline of 7^th^ cervical spinal cord segment (tissue displacement = 0.800mm). The SCI was performed using a Louisville Injury System Apparatus (LISA) device according to our previously established protocol. After injury, the muscles were sutured in layers, and the skin was closed with clips. The rats were allowed to recover in a warmed cage with water and food easily accessible. The analgesic buprenorphine (0.05-2.0 mg/kg) was subcutaneously delivered every 6-12 h post-surgery for 2 d. For post-surgical care, bladder expression was performed 3 times a day.

### Spinal vascular imaging in vivo

The details of this procedure can be found in our previous publication. Briefly, while the rat is under deep anesthesia, a specialized catheter connected with a 1 ml saline-filled syringe is inserted and secured via the external jugular vein. A laminectomy from the 5^th^ to 7^th^ cervical segments was performed after vertebral stabilization, followed by sealing the surrounding muscle-bone field with tissue adhesive. A line of 4% agar was built around the edge to form the wall for a two-photon imaging window. For imaging, the rats were imaged using a two-photon laser-scanning microscope (TPLSM, Prairie Technologies, Middleton, WI). Excitation was generated by a tunable Maitai Ti:sapphire laser (Newport, Mountain View, CA) tuned to 860 nm. Band-pass filtered fluorescence (560–660 nm) was collected by photomultiplier tubes of the system. The same fields were identified and imaged at all different time points. For pre-injury imaging, the rat was placed under the microscope focused on the exposed spinal segments (C5-C7) and 0.5 mL of Rhodamine B isothiocyanate-dextran (Rho-dextran, 70kDa, Sigma-Aldrich, 4 mg/mL, dissolved in saline) was injected slowly via the aforementioned catheter. At 4x magnification (NA 0.1; Olympus, Shinjuku, Tokyo, Japan), a bright-field image of the surface blood vessel pattern was acquired as a location map using a charge coupled device (CCD) camera. At 10x magnification (NA 0.30; Olympus), z-stack images across C5-C7 segments (epicenter, transitional zone, remote zone, 200-400 µm thickness) were taken to track dynamic changes in spinal blood vessels, including vessel diameter and vascular permeability. For post-injury time points, 0.5 mL of Fluorescein isothiocyanate-dextran (FITC-dextran, 70kDa, 4 mg/mL, dissolved in saline) was slowly injected. The same imaging steps were performed as the pre-injury imaging. Later the Z-stack images were analyzed to generate changes in vessel diameter and vascular permeability using imageJ.

In the first study, a total of 32 rats in the *in vivo* spinal vascular imaging experiment were imaged at different acute time points following contusion, 8 in each of the Sham, 0.5 hour post-injury (hpi), 2 hpi and 4 hpi groups. These animals were terminated after imaging (terminal experiments). In the second study, there were two groups in this experiment: FA-GC treated group (*n* =11) and saline control group (*n* =7).

### Measurement of vessel diameter and vascular permeability change

The acquired Z-stack images were analyzed via ImageJ to measure changes in vessel diameter and vascular permeability. The images were first calibrated true to scale. For vessel diameter, a line is drawn perpendicular to the long axis of the vessel segment to measure the diameter. The diameter of the blood vessel is determined by the average of three measurements across the vessel segment. For vascular permeability, a probe of a 200 μm line is drawn both inside and outside the vessel to measure the fluorescence integrated density. The vascular permeability is calculated as the ratio of the outside value divided by the inside value.

### Preparation and delivery of FA-GC nanoparticles in vivo

Ferulic acid–glycol chitosan nanoparticles (FA-GC) were dissolved in 0.9% sodium chloride solution by sufficient sonication to produce a homogeneous solution of FA-GC nanoparticles. Each aliquot was 1 ml and prepared at the day of surgery. FA-GC (22.7mg/kg, 1ml in saline) were intravenously administrated to the SCI rats via the same EJV catheter previously used for tracer injection within 15 min post contusive injury.

### Tissue processing

For the groups of animals assigned for histological analysis, tissues were collected and processed using methods adapted from a previous publication. Anesthetized animals were transcardially perfused with 100 ml 0.01M phosphate buffer saline, followed by 400 ml of ice-cold 4% paraformaldehyde fixative solution in 0.01M PBS. A 9 mm segment of cervical cord including C5 to C7 was collected and post-fixed in the same fixative solution overnight, and then transferred to a solution of 30% sucrose in 1x PBS for 5 d. Spinal cord segments were embedded in O.C.T compound (Fisher, Waltham, MA) for cryostat sectioning. The tissues were cut on a cryostat (LeicaCM 1950; Buffalo Grove, IL) in a series of coronal 25-µm-thick sections from C5 to C7 mounted on slides, and stored at –20°C. The sections were then processed for immunofluorescence labeling.

In the first study, a total of 44 rats in the histological experiment were divided at different acute time points following contusion, 8 in each of the Sham, 0.5 hpi, 2 hpi and 4 hpi groups, and 2 additional groups with 6 in each of the 1 and 7 day post-injury (dpi) groups. In the second study, there were two groups in this experiment: FA-GC treated group (*n* = 8) and saline control group (*n* = 8).

### Immunohistochemistry and immunocytochemistry

The spinal cord sections were washed in 1xPBS 3 times at room temperature, blocked in 10% normal goat serum (NGS) or normal donkey serum (NDS) blocking solution (with 0.1% Triton X-100 in 0.01M PBS) for 1 h, then incubated with primary antibodies in 5% NGS or NDS and Triton X-100 (0.05% in 0.01M PBS) at 4°C overnight. After washing in 1xPBS 3 times, sections were incubated with host-specific secondary antibodies for 1 h at room temperature, followed by DAPI staining for 5 min. The sections were washed in 1xPBS again 3 times and coverslips were mounted with Fluoromount-G (SouthernBiotech, Birmingham, AL). The primary antibodies and their final dilutions were used as follows: anti-RECA1 (blood vessels; 1:200, mouse; Bio-Rad, Hercules, CA); anti-GFAP (astrocytes; 1:1000, chicken; Abcam, Cambridge, United Kingdom); anti-APC (oligodendrocytes; 1:400, mouse; Abcam); anti-Iba1 (microglia; 1:500, goat; Abcam); anti-laminin (basement membrane of blood vessels; 1:200, rabbit; Millipore); anti-NeuN (neurons; 1:1000, mouse or rabbit; Millipore); anti-SMI31 (axons; 1:1000, mouse; Biolegend, San Diego, CA); anti-MBP (myelin; 1:200, rat; Millipore). Secondary antibodies with either goat or donkey host were used in a dilution of 1:1000 (details are shown in **Table 1**).

**Table 1.**
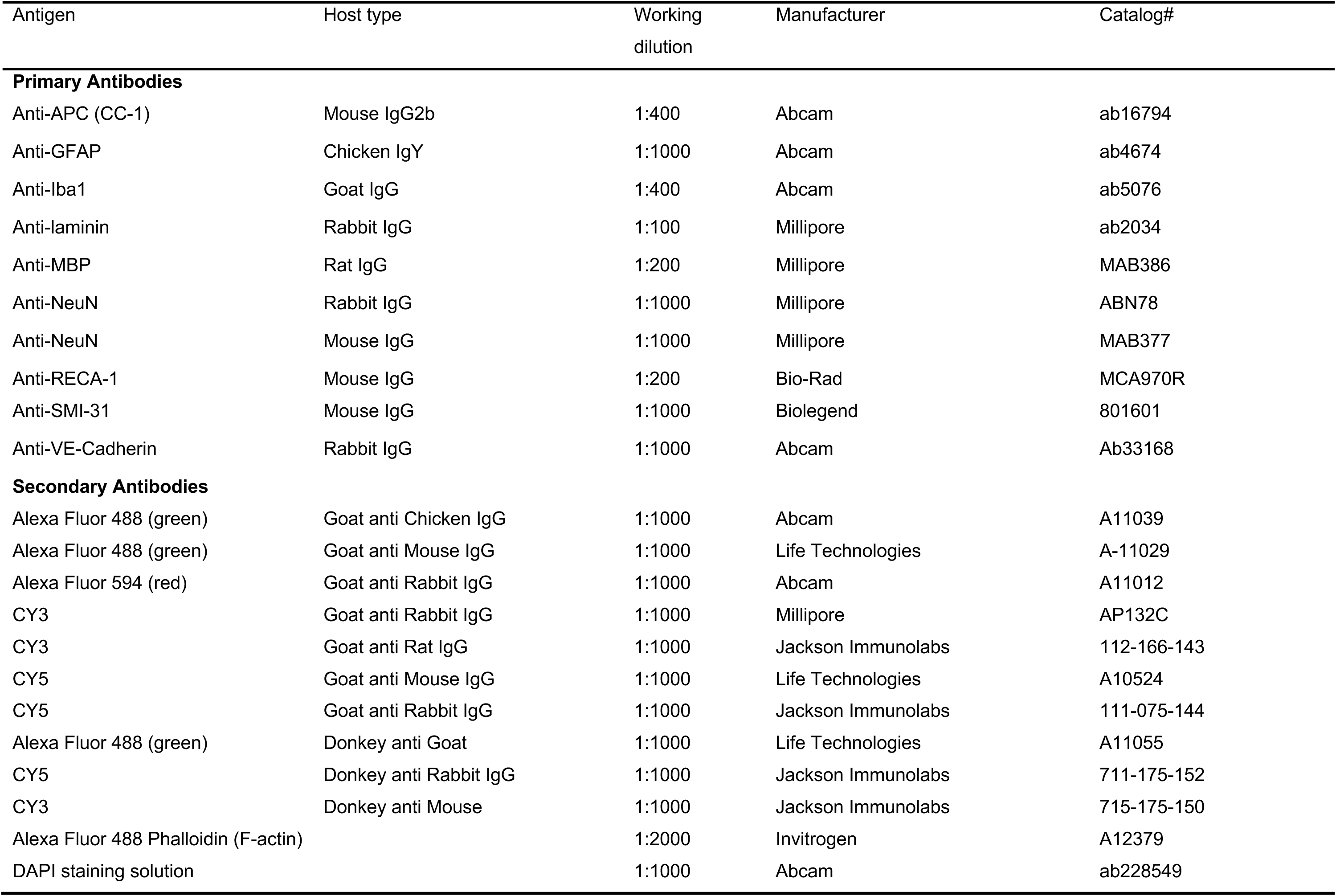
Primary and secondary antibodies used in this study.

**Table 2.** Spatiotemporal cellular changes following SCI.

|  | Remote region (C5) | Transitional region (C6) | Epicenter (C7) |
| --- | --- | --- | --- |
| Blood vessel | • Vessel diameter remains the same until a much later time.<br>• Vessel leakage remains the same in the ventral gray matter; some IgG positive vessels occur in the dorsal column at 1 d. | • Vessel diameter is bigger.<br>• Vessel leakage happens at the same time with epicenter but smaller range. | • Vessel diameter is bigger.<br>• Vessel leakage (hemorrhage & plasma leaking) is instantly there. |
| Neuron | • Neuron density remains the same. | • Neuron density is smaller starting at 1 d after injury. | • Neuron density is smaller starting at 4 h after injury. |
| Astrocyte | • Cell number remains the same.<br>• GFAP expression remains the same. | • Cell number remains the same.<br>• GFAP expression increases at 4 h. | • Cell number first becomes smaller at 4 h but increases at 7d.<br>• GFAP expression increases at 2 h. |
| Microglia | • The percentage of hypertrophied over ramified cells increases at 1 d.<br>• The total microglia number increases at 7 d. | • The percentage of hypertrophied over ramified cells increases at 1 d.<br>• The total microglia number increases at 7 d. | • The percentage of hypertrophied over ramified cells increases at 2 h.<br>• The total microglia number increases at 7 d. |
| Axon | • Axon number is reduced later (1 d). | • Axon number is reduced later (1 d). | • Axon number is reduced instantly (0.5 h after injury). |
| Myelin | • Same as epicenter | • Same as epicenter | • Myelin area become bigger because of dissociation from wrapping the axons, then smaller because of disassembly. |
| Oligodendrocyte | • Cell number decreases starting at 1 d. | • Cell number decreases starting at 1 d. | • Cell number decreases in the dorsal column starting at 0.5h. |

Abbreviations: GFAP, glial fibrillary acidic protein; IgG, immunoglobulin G; SCI, spinal cord injury.

For co-labeling with IgG and laminin, sections were stained with primary and secondary antibodies against laminin regularly as mentioned above; during the step of secondary antibody incubation, a fluorescence-tagged anti-rat IgG antibody (Cy3, 1:1000; Jackson ImmunoResearch Laboratories, Inc., West Grove, PA) was added for 1 h at room temperature.

For immunocytochemistry, HBMECs grown on collagen-coated coverslips were fixed with 4% paraformaldehyde followed by blocking with goat serum in 0.01M PBS for 1 hour at room temperature. The cells were incubated with rabbit anti-VE-Cadherin (1:200; Abcam) primary antibody overnight at 4°C. After rinses in PBS, cells were then incubated with goat secondary antibodies conjugated with Alexa 488 or Cy3 (1:1000, Jackson ImmunoResearch Laboratories, Inc.). F-actin staining was accomplished by Alexa Flour 488-conjugated phalloidin (1:2000; Invitrogen). For nuclear labeling, cells were then counterstained with DAPI. After final PBS rinses, coverslips were mounted on glass slides with Fluoromount-G (SouthernBiotech, Birmingham, AL). Fluorescence images were captured using an inverted microscopy system (Zeiss Axiovert 200M, Jena, Germany) combined with an apotome and a digital camera (Zeiss Axio Cam MRc5, Jena, Germany). Immunofluorescence intensity of VE-Cadherin and F-actin was quantified by ImageJ.

### Microscopy

For immunohistochemistry, an inverted fluorescent microscopy system (Zeiss Axiovert 200 M, Jena, Germany) was used to capture fluorescent images. The microscope was combined with an apotome and interfaced with a digital camera (Zeiss Axio Cam MRc5) controlled by a computer. Images were captured using the apotome with software (AxioVision v4.8) and output as a cut view image. The images were later assembled and quantified in Photoshop CC (Adobe System, San Jose, CA).

### Cell counting

At each time point, cells were counted in one stacked image from each region: C5 (remote segment), C6 (transitional segment), and C7 (epicenter) (25 μm thickness, every 120^th^ section from the epicenter, 3000 μm apart, based on the topographic info of the rat spinal cord, 3 sections for each animal). Every single positive cell (even partial positive cell at the borders of sections) was counted in multi-planes throughout the entire designated area in the 25 μm section.

To quantify leaked vessels, the sections were processed with antibodies against IgG and laminin. Both dorsal white matter and ventral gray matter were examined. The percentage of leaked vessels were calculated by the leaked vessel number over total vessel number. To quantify the number of surviving neurons, the sections were processed with antibody against NeuN and ventral gray matter was examined. Cell density was determined by dividing the total NeuN-positive number by the area of contoured ventral gray matter, expressed as average cell number per mm^2^. In the second study, the spinal cord sections at 7 dpi (n = 8 for both groups; FA-GC treatment and saline control groups) were double labeled with primary antibody against NeuN and DAPI, followed by secondary antibodies. One side of the gray matter in the spinal cord was examined. Cell density was then calculated by dividing the total NeuN-positive number by the area of gray matter, and the average cell number per mm^2^ was reported as total neuronal density. The gray matter area was further divided into three sub-regions based on the ten-layer Rexed laminae system: dorsal horn (lam I-VI), intermediate zone (lam VII), and ventral horn (lam VIII-IX). The neuron density in each sub-region was calculated the same as total neuronal density. To quantify microglia, the total number and percentage of ramified and hypotrophy cells were examined in the ventral gray matter using antibody against Iba1; similar as above, cell density was expressed as average Iba1^+^ number per mm^2^ and the percentage of ramified and hypotrophy microglia were calculated by the ramified or hypotrophy cell number over the total number; to quantify the area of microglial process, a 300 μm x 300 μm area was selected in the ventral gray matter, 500 μm away from the midline and the area in mm^2^ was measured using ImageJ. To quantify astrocytes, antibody against GFAP was used and the cell density in the ventral gray matter was expressed as average number per mm^2^. Same as microglia, the same fixed area was selected and the average astrocyte area in mm^2^ (Total GFAP area divided by cell number) and integrated density of GFAP were measured using ImageJ. To quantify axons and myelination, antibodies against SMI-31 and MBP were used and the designated dorsal white matter area was examined; both axon and myelin were measured using ImageJ, expressed as total axon count, average axon size in mm^2^, total myelin area, and myelin area per axon in mm^2^. To quantify the number oligodendrocyte in the dorsal white matter, antibody against APC (CC-1) was used and cell density was calculated the same way as described above.

### Forelimb functional assessment: Modified Grip Strength

A modified forelimb grip strength test was applied on a grip strength meter (TSE Systems, Inc., Chesterfield, MO) to assess skeletal muscle function according to a previous publication. A square metal frame with horizontal metal bars was connected to a high-precision force sensor. The metal frame was stably fixed on the metal stand vertically and the parallel metal bars were horizontally placed above the ground with a fixed height. Before each trial, the gauge was reset to 0. After placing the animal to grasp the same horizontal bar, the rat’s tail was pulled by an examiner in a downward direction. As the rat pulls and releases the bar, the maximum force exerted was shown on the digital display of the force sensor as unit of pond (1 pond (P) = 0.0098 newton (N)). The criteria to validate a trial is that trials were excluded if only one forepaw, or the hindlimbs were involved; or the rat turned or left the bar without resistance during the test. At one-minute intervals, 10 consecutive measurements were performed on individuals and the average was reported as the grip strength. Given that the measurement can be influenced by the speed of the tail pull, the procedure was performed at a constant speed sufficiently slow to allow the rat to build up a resistance against it. The rat’s body weight was measured after the test. To check if changes in body weight have a major effect on grip strength results, a Pearson correlation coefficient was analyzed between the body weight and the grip strength.

### In vitro BBB model

Primary Human Brain Microvascular Endothelial Cells (HBMECs) were purchased from Cell Systems (ACBRI 376, Kirkland, WA, USA). HBMECs were cultured in Clonetics EGM-2 MV media (CC-3202, Lonza, Walkersville, MD, USA), and only up to nine passages were used for analyses. The *in vitro* BBB model was established in permeable transwell inserts as previously described. The transwell PET membranes (0.4-μm pore, 11-mm diameter; Corning, Lowell, MA, USA) were pre-coated with collagen (15 μg ml^-1^) and fibronectin (30 μg ml^-1^) before cell culture. HBMECs were placed onto the membrane at a density of 2.5 x 10^5^ cells per membrane. Cultures were maintained at 37 °C in a cell culture incubator (95% air and 5% CO_2_) for 4 days to reach monolayer confluence. To assess the permeability of the cell layer, 4.4 kDa TRITC-dextran or 70 kDa FITC-dextran (Sigma-Aldrich) were added into the donor chamber at a concentration of 1 mg ml^-1^ in 550 μl media. A sample of 50 μl media was removed from the receiving chamber at different time points. Fluorescence intensity was measured by with a fluorescence plate reader at baseline, 1 h, 3 h, 6 h, 18 h, and 24 h. The concentrations of tracers were calculated based on a standard curve fitted using known concentrations of tracers. Fifty microliter fresh media was added back after each reading. BBB permeability was calculated by measuring the diffusion coefficient of tracers from the luminal to the abluminal chamber.

### Oxygen Glucose Deprivation (OGD)

Cultured HBMECs were exposed to OGD for 120 min. After the medium was replaced with glucose-free medium, and cultures were placed inside a Billups-Rothenberg modular incubator chamber (Del Mar, San Diego, CA, USA), which was flushed with 100% nitrogen for 5 min and then sealed. The entire chamber was then placed in a water-jacketed incubator (Forma, Thermom Fisher Scientific, Waltham, MA, USA) at 37 °C for 120 min and then returned to 95% air, 5% CO_2_ and replaced with glucose-containing culture medium. As a control, cultures with glucose-containing medium were incubated at 37 °C in humidified 95% air and 5% CO_2_ during the same period of time. The FA-GC was added immediately after OGD and remained in the media after OGD.

### Statistical analysis

Both *in vivo* diameter and cell quantification data were presented as average ± standard deviation and analyzed using a one-way ANOVA of variance test followed by a Tukey post hoc test using GraphPad Prism 7 (GraphPad Software, Inc.; La Jolla, CA). Statistical significance was set at *p* < 0.05. *In vivo* permeability data were analyzed using repeated measure two-way ANOVA followed by LSD as post-hoc test via SPSS software (IBM Corporation, Armonk, NY). Significance was set at *p* < 0.05.

The data of *in vivo* diameter were analyzed using a one-way ANOVA of variance test followed by a Dunnett’s multiple comparison test. The quantification of the neuron was presented as average ± standard deviation and analyzed using a one-way ANOVA of variance test followed by a Tukey post-hoc test. Behavior data were presented as average ± standard deviation and analyzed using unpaired t test at different time points. All analyses above are using GraphPad Prism 7 (GraphPad Software, Inc.; La Jolla, CA). *In vivo* permeability data were analyzed using repeated measure two-way ANOVA followed by LSD as post-hoc test via SPSS software (IBM Corporation, Armonk, NY). All significances were set at *p* < 0.05.

## Acknowledgments

FA-GC nanoparticles were provided by Dr. Ji-Xin Cheng’s laboratory. We thank Patti L. Raley, a Medical Editor, for her help with reviewing and editing the manuscript and Dr. Xingjie Ping for constructive suggestions. We also thank Mr. Christopher M. Brown for his excellent illustrative work.

## Author contributions

C.C. designed and performed the experiments, and analyzed the data; Y.S. performed immunostaining, imaging capture, and provided technical assistance with cell culture; XL.D. and H.D. performed behavior assessments and provide assistance for animal surgery; XT.W. and Q.H. provided constructive suggestions and revised the manuscript; WH.X. and XM.J. provided technical assistance with *in vivo* imaging; ZY.P and C.B.S provided technical assistance for the SCI models; S.Y.L. and J-X.C. provided the nanoparticles; W.W. is the corresponding author who conceived, designed, and directed the project; C.C. and W.W. wrote the manuscript.

## Data and materials availability

All data are available in the main text or the Supplementary Materials; raw imaging data and analysis code are available to editors and reviewers via private links and will be made public upon publication. FA-GC is available under an MTA.

## Supplemental Figure Legends

**Supplemental Figure 1.**
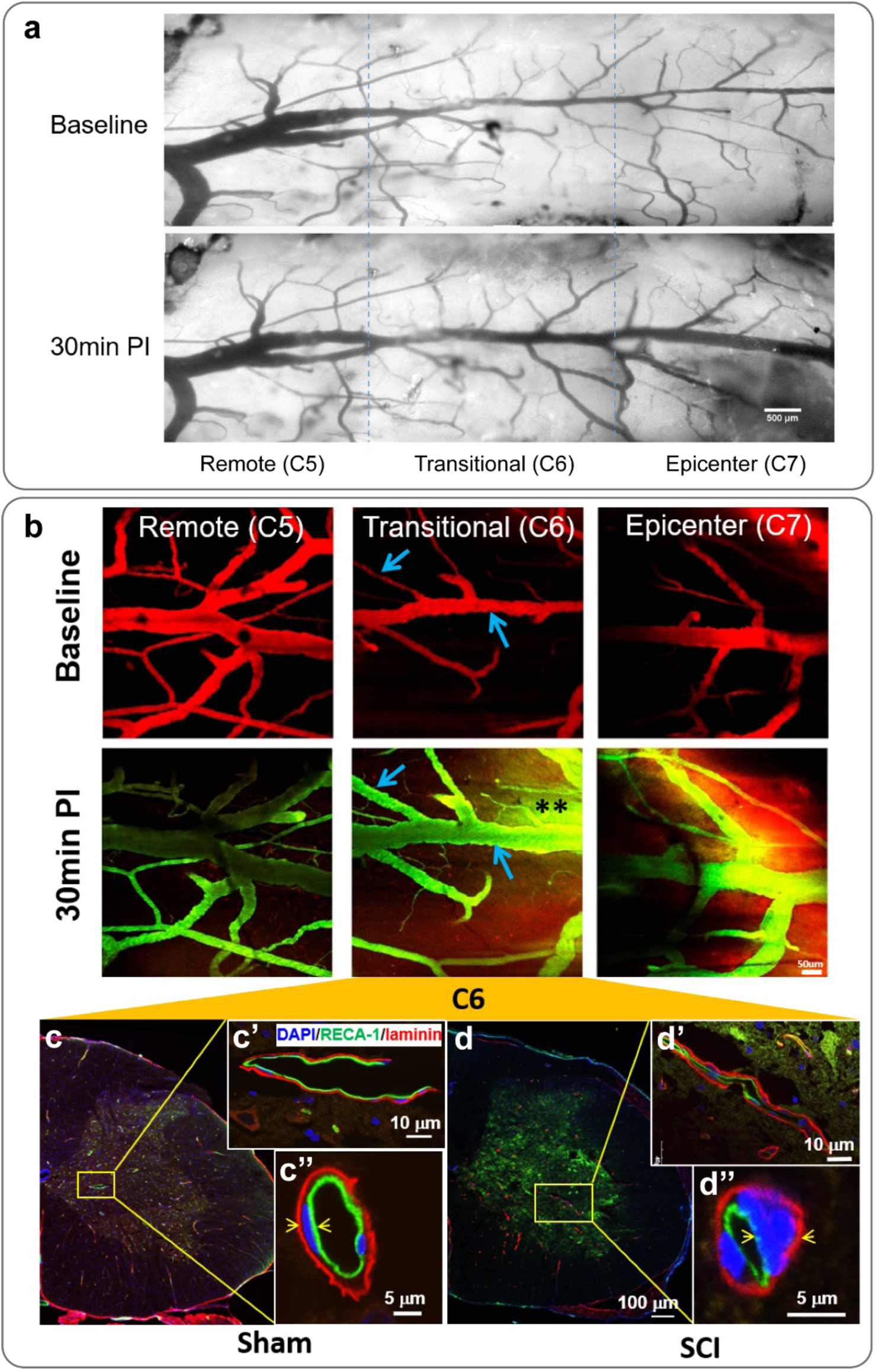
Evidence for vascular dysfunction after a C7 midline contusive SCI. **(a)** The diameter of blood vessels were increased in the epicenter (C7) and transitional (C6) segments at 30 minutes (min) after a C7 midline contusive spinal cord injury (SCI). **(b)** At 30 min post-injury (PI), the vessel diameter and vascular permeability were both visually increased in the epicenter and transitional segments comparing to baseline. Blue arrows indicate measured blood vessels, and ** indicates the leaking area. **(c-d)** In the transitional segment, the spinal cord cross-sections of sham or SCI were stained with RECA-1 (green) for endothelial cell layer and laminin (red) for basement membrane, and counterstained with DAPI (blue) for nuclear staining. **(c’, c’’, d’, d’’)** Enlarged areas display the staining in different views. The SCI group showed enlarged nucleus whereas the nucleus in the sham group was in a slim and elongated shape, indicated by pairs of yellow arrows.

**Supplemental Figure 2.**
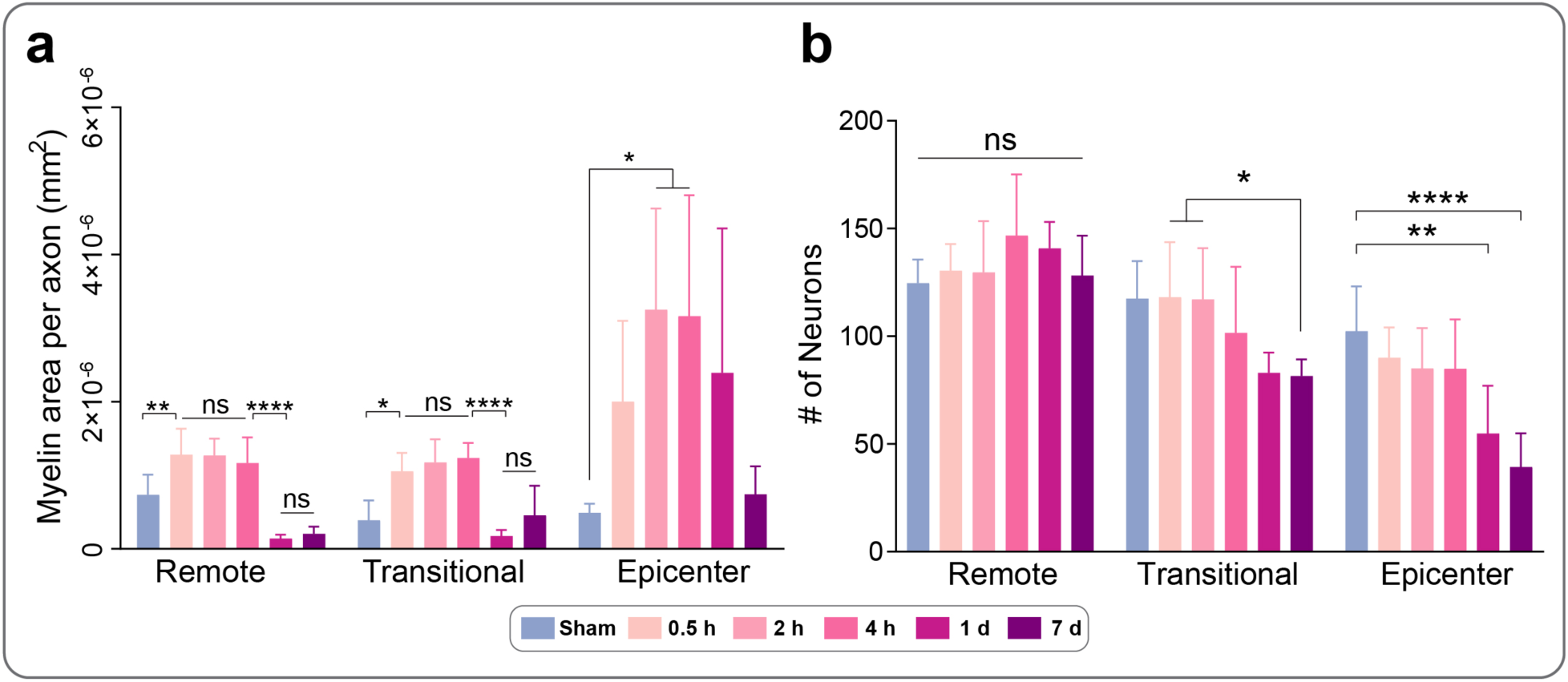
Spatiotemporal axon/myelin and neuronal changes after a C7 midline contusive SCI. **(a)** the myelin area per axon in the dorsal column, and **(b)** the number of neurons in the ventral gray matter were quantified. For **(a)** and **(b)**, n = 7-8 for sham, 0.5 hour (h), 2 h, and 4 h post-injury. n = 6 for 1 d and 7 d post-injury. For each segment, one-way ANOVA followed by Tukey’s multiple comparison tests were performed. ns, not significant; * P < 0.05; ** P < 0.01; *** P < 0.001; **** P <0.0001. All data were presented as mean ± s.d.

**Supplemental Figure 3.**
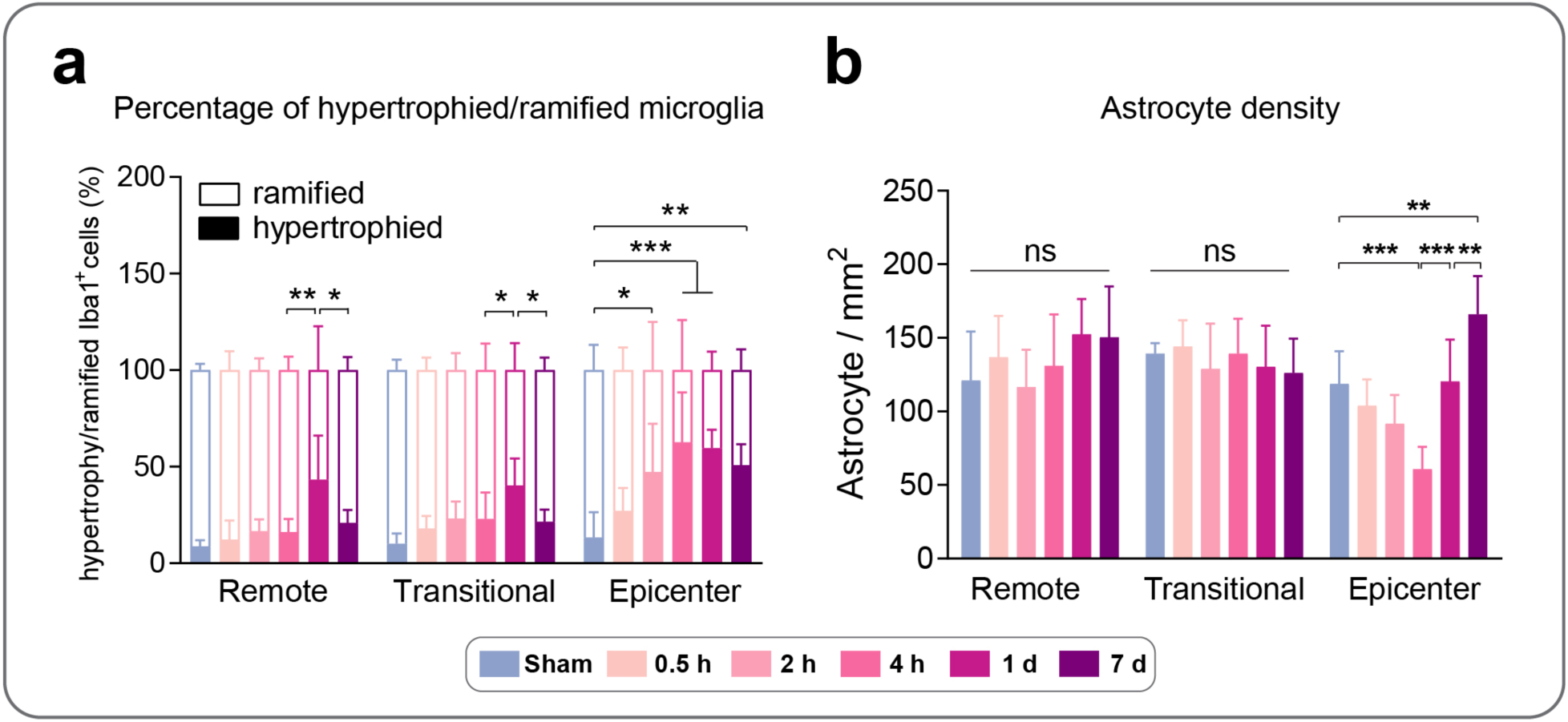
Spatiotemporal glial cell changes in the ventral gray matter after the C7 midline contusive SCI. **(a)** The percentage of hypertrophied and ramified microglia (n = 6 per group), and **(b)** the cell density of astrocytes in the ventral gray matter (n = 7-8 for sham, 0.5 hour (h), 2 h, and 4 h post-injury. n = 6 for 1 d and 7 d post-injury) were quantified. For each segment, one-way ANOVA followed by Tukey’s multiple comparison tests were performed. ns, not significant; * P < 0.05; ** P < 0.01; *** P < 0.001; **** P <0.0001. All data were presented as mean ± s.d.

**Supplemental Figure 4.**
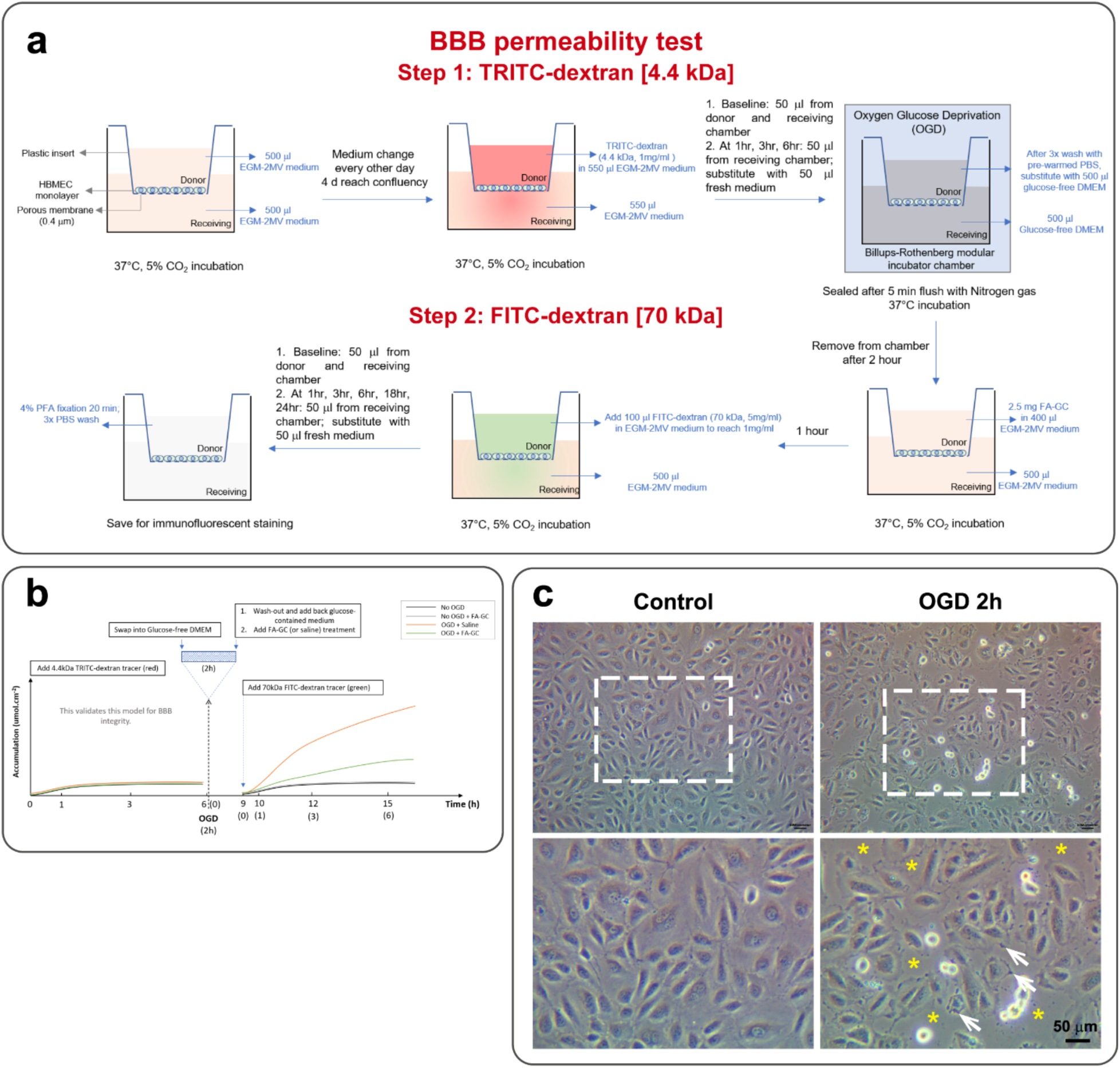
*In vitro* BBB model and permeability test. **(a)** Illustration of details in the BBB permeability test in two consecutive steps. **(b)** The timeline of the permeability test. **(c)** Brightfield images of human brain microvascular endothelial cells (HBMECs) monolayer cell culture were taken after oxygen-glucose deprivation (OGD) (2-hour [h] duration) or control conditions. White dashed lines framed enlarged areas. White arrows indicate the edges of endothelial cells, and yellow * indicates gaps between endothelial cells. After 2-h deprivation of oxygen and glucose, the integrity of HBMEC monolayer was disrupted, represented by larger gaps between cells (yellow *).

**Supplemental Figure 5.**
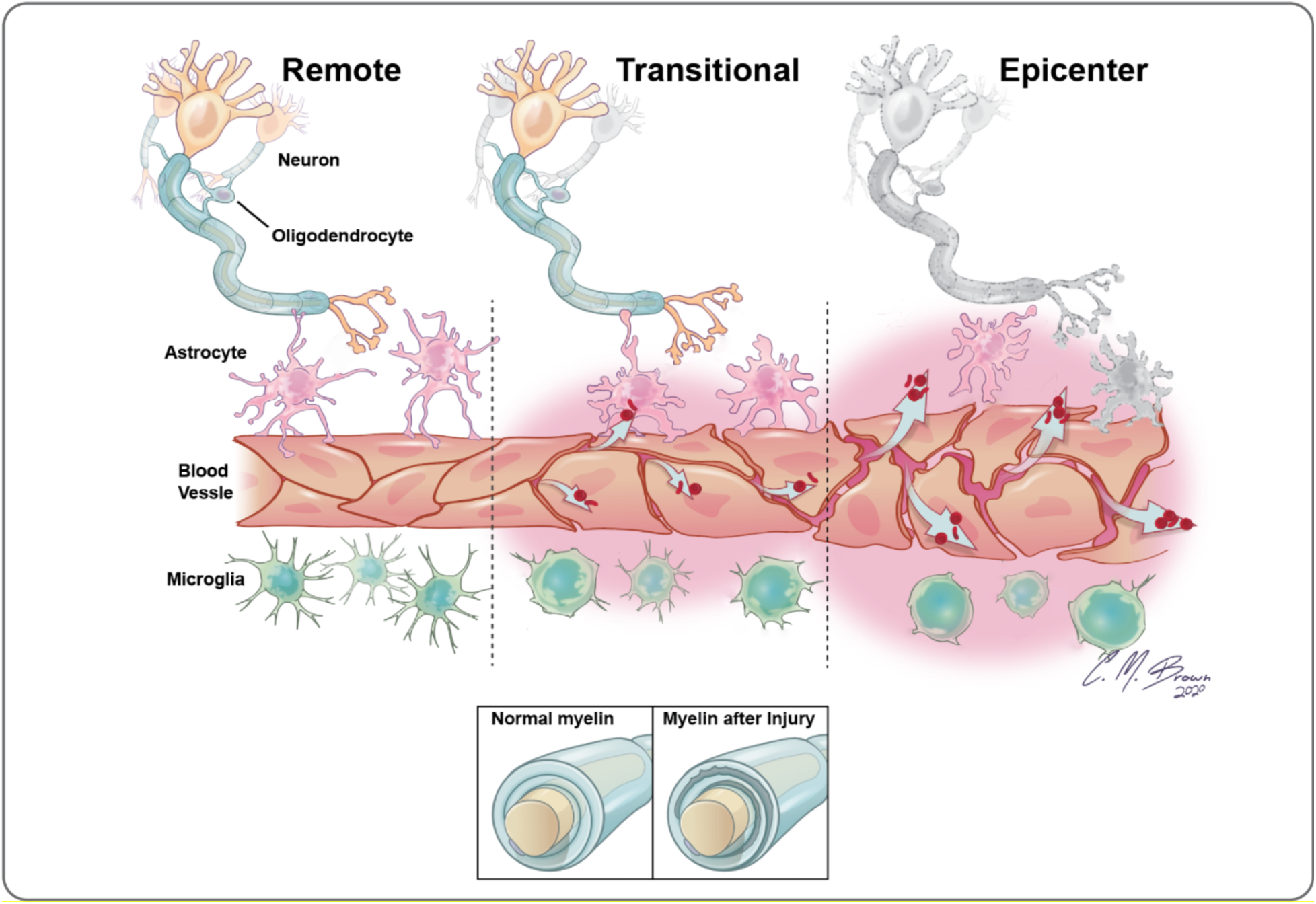
Schematic summary. The schematic diagram depicts the acute region-specific cellular changes following a contusive spinal cord injury (SCI). Vascular leak and vasodilation were first detected in the epicenter, followed by a cascade of secondary neuronal loss and glial responses within hours. While acute vascular disruption also occurred simultaneously in the transitional region, a delay between vascular leak and loss of neuronal components was found. The remote region remained relatively normal with least pathological changes.

